# The crosstalk between DNA-PK and cGAS drives tumor immunogenicity

**DOI:** 10.1101/2022.06.08.495278

**Authors:** Clara Taffoni, Johanna Marines, Hanane Chamma, Mathilde Saccas, Amel Bouzid, Soumyabrata Guha, Ana-Luiza Chaves Valadao, Katarzyna Polak, Maguy Del Rio, Celine Gongora, Donovan Pineau, Jean-Philippe Hugnot, Karima Kissa, Laura Fontenille, Fabien P. Blanchet, Isabelle K. Vila, Nadine Laguette

## Abstract

Cytosolic DNAs promote inflammatory responses upon detection by the cyclic GMP-AMP (cGAMP) synthase (cGAS). It has been thus suggested that cGAS downregulation is an immune escape strategy harnessed by tumor cells. Here, we used glioblastoma cells that lack cGAS to question whether alternative DNA detection pathways can promote pro-inflammatory signaling. We show that the DNA-PK DNA repair complex drives cGAS independent inflammatory responses but that its catalytic activity is required for cGAS-dependent cGAMP production and optimal downstream signaling. We further show that the cooperation between DNA-PK and cGAS favors the expression of chemokines that promote macrophage recruitment in the tumor microenvironment, a process that impaired early tumorigenesis but correlated with poor outcome. Thus, our study supports that cGAS-dependent signaling is acquired during tumorigenesis and that cGAS and DNA-PK activities should be analyzed concertedly to predict the impact of strategies aiming to boost tumor immunogenicity.

## INTRODUCTION

Most cells mount type I Interferon (IFN) responses in the presence of cytosolic DNA ^1^. One of the key pathways involved in the detection of immune-stimulatory DNA relies on the cyclic GMP-AMP synthetase (cGAS). Upon detection of dsDNA, ssDNA or RNA:DNA hybrids ^2, 3^, cGAS produces the cyclic GMP-AMP (cGAMP) second messenger. Interaction of cGAMP with the Stimulator of Interferon Genes (STING) promotes the assembly of a signalosome comprised of the Tank Binding Kinase 1 (TBK1) and transcription factors such as Interferon Regulatory Factor 3 (IRF3) ^4^. TBK1-dependent phosphorylation of IRF3 leads to its nuclear translocation and subsequent activation of transcriptional programs that ultimately lead to the production of inflammatory cytokines, chemokines and type I IFNs ^5^.

The cGAS-STING signaling cascade has been shown to be essential to the orchestration of antitumor responses ^6, 7^. Indeed, activating the cGAS-STING axis can promote tumor rejection through increasing tumor immunogenicity and priming T cell responses ^8, 9^. However, STING activation can also foster metastatic dissemination ^10^, and impair the establishment of durable immunity ^11^. There is no clear mechanistic explanation for these discrepancies, although the diversity of cells composing the tumor microenvironment, and their differential expression of cGAS and/or STING may be a key parameter ^12^. Downregulation of the cGAS-STING axis has therefore been proposed to be an immune escape strategy exploited by tumor cells ^13, 14^, despite evidence that high expression of cGAS and/or STING predicts poor outcome for cancer patients ^15^.

Recently, the DNA-dependent protein kinase (DNA-PK) complex, involved in the repair of double strand DNA lesions by non-homologous end-joining (NHEJ) ^16^, has been involved in the detection of DNA virus-derived cytosolic dsDNA, eliciting type I or III IFN responses ^17–21^. The DNA-PK core complex is a holoenzyme comprised of the KU70*^XRCC^*^6^ and KU80*^XRCC5^* subunits that ensure the recruitment of the DNA-dependent protein kinase catalytic subunit (DNA-PKcs*^PRKDC^*) to double strand breaks ^22^. How cGAS- and DNA-PK-mediated detection of dsDNA are coordinated remains the subject of controversies. Indeed, there are reports indicating that DNA-PKcs inhibits cGAS ^23, 24^, while others suggest that DNA-PK may be required for cGAS-STING-dependent inflammatory responses to viral DNA ^17^. Intriguingly, despite the crucial role of DNA-PK in NHEJ, and the tight link between DNA repair machineries and nucleic acid sensing ^25^, there is no evidence for a role of DNA-PK in eliciting inflammatory responses following genotoxic stress ^19^.

Here, we used glioblastoma cells to interrogate how type I IFN responses are initiated in the absence of cGAS. We found that DNA-PK can promote nucleic acid- and chemotherapy-associated inflammatory responses independently of cGAS. Further, we uncover that cGAS and DNA-PK cooperate for optimal STING-dependent signaling, thereby defining tumor immunogenicity. Our work thus suggests that cGAS-dependent signaling is acquired during tumorigenesis.

## RESULTS

### DNA-PK catalytic activity promotes cytosolic dsDNA-dependent type I Interferon responses in cGAS-deficient cells

Assessment of dsDNA-induced type I IFN responses in the T98G glioblastoma cell line, that does not express cGAS (Figure 1A and S1A), showed increased Interferon β (*IFNB*) and C-X-C motif chemokine ligand 10 (*CXCL10*) Interferon stimulated gene (ISG) mRNA levels (Figure 1B), attesting to the activation of type I IFN responses. Similar analysis of type I IFN responses, conducted on CD133+ glioblastoma stem cells (namely, Gli4 and Gli7) ^26^, showed that despite the absence of cGAS (Figure 1C and S1B), these cells are also able to express *IFNB* and *CXCL10* (Figure 1D and S1C) upon challenge with dsDNA. Western blot (WB) analysis further showed that dsDNA stimulation of T98G, Gli4 and Gli7, led to phosphorylation of IRF3 (pIRF3) (Figure 1A, C and S1B). This indicates that glioblastoma cells possess cGAS-independent cytosolic dsDNA detection mechanisms.

**Figure 1:**
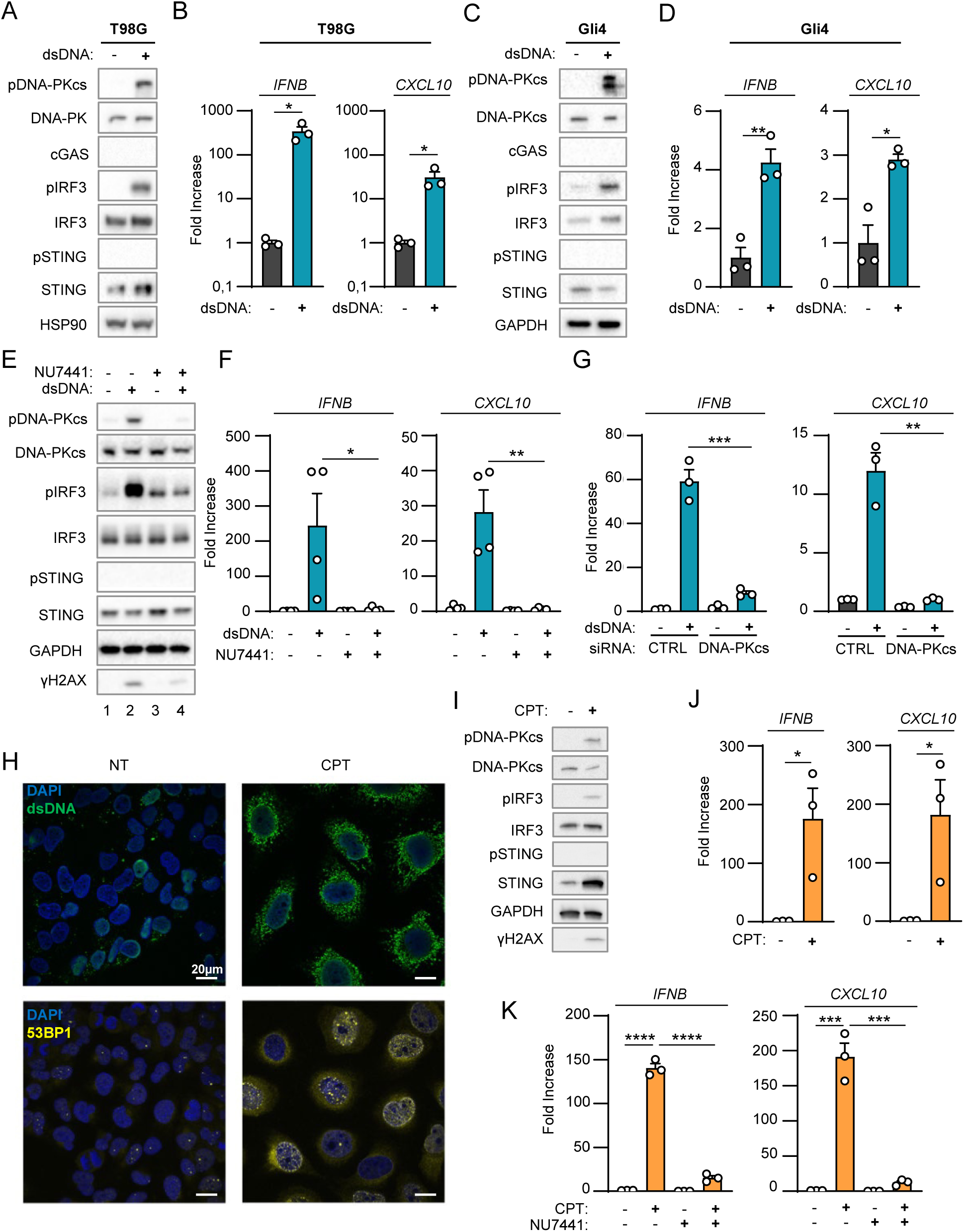
DNA-PK catalytic activity promotes nucleic acid-dependent type I IFN responses in cGAS-deficient cells. (A) T98G cells were challenged or not with dsDNAs for 6 h prior to whole cell extraction and Western Blot (WB) analysis using indicated antibodies. (B) *IFNB* and *CXCL10* mRNA levels were analyzed by RT-qPCR in samples treated as in A (n=3). (C) Gli4 cells were treated as in A prior to WB analysis using indicated antibodies. (D) *IFNB* and *CXCL10* mRNA levels were analyzed by RT-qPCR in samples treated as in C (n=3). (E) T98G cells were challenged or not with dsDNA for 6 h, in the presence or not of the NU7441 DNA-PKcs inhibitor, prior WB analysis using indicated antibodies. (F) *IFNB* and *CXCL10* mRNA levels were analyzed by RT-qPCR in samples treated as in E (n=4). (G) T98G cells were treated with non-targeting (CTRL) or DNA-PKcs-targeting siRNAs for 72 h prior to 6 h challenge with dsDNA. *IFNB* and *CXCL10* mRNA levels were analyzed by RT-qPCR. Graphs present a representative biological triplicate (n=3). (H) T98G cells were treated or not with 0.16 µM camptothecin (CPT) for 48 h prior to immunofluorescence analysis using dsDNA- and 53BP1-specific antibodies, and DAPI nuclear staining. (I) Whole cell extracts from T98G cells treated or not for 72 h with 0.16 µM CPT were analyzed by WB using indicated antibodies. (J) *IFNB* and *CXCL10* mRNA levels were analyzed by RT-qPCR in samples treated as in I (n=3). (K) T98G cells were treated or not with 0.16 µM CPT for 72 h, in presence or not of NU7441, prior to assessment of *IFNB* and *CXCL10* mRNA levels by RT-qPCR. Graphs present a representative biological triplicate (n=3). All immunoblots are representative experiments. All graphs present means ± standard error from the mean (SEM). P-values were determined by Student’s t-test. ns: not significant. * p < 0.05, ** p < 0.01, *** p < 0.001, **** p < 0.0001.

Because the DNA-PK complex was previously reported as an alternative cytosolic dsDNA sensor, we interrogated whether DNA-PK could be responsible for the type I IFN response elicited by dsDNA in absence of cGAS. To this aim, we first performed WB analysis using an antibody specific for the auto-phosphorylation of DNA-PKcs on Serine 2056 (pDNA-PKcs), which reflects its activation ^27^. Stimulation of T98G, Gli4 and Gli7 with dsDNA led to DNA-PKcs phosphorylation (Figure 1A, 1C and S1B). Second, we tested whether DNA-PK is responsible for type I IFN responses in absence of cGAS in glioblastoma cells by treating T98G and Gli4 with the NU7441 DNA-PKcs inhibitor). WB analysis showed that treatment with NU7441 inhibited dsDNA-induced DNA-PK auto-phosphorylation and decreased phosphorylation of the H2AX DNA-PK substrate (Figure 1E, compare lanes 2 and 4), attesting to efficient DNA-PKcs inhibition. NU7441 treatment also led to a decrease of pIRF3 levels (Figure 1E) and of *IFNB* and *CXCL10* levels (Figure 1F and S1D). Finally, T98G cells were treated with scrambled or DNA-PKcs-targeting siRNAs prior to analysis of dsDNA-dependent type I IFN responses. Knock-down of DNA-PKcs in T98G cells (Figure S1E) abrogated dsDNA-induced type I IFN responses (Figure 1G). Thus, DNA-PKcs drives dsDNA-induced type I IFN responses in cells lacking cGAS.

### DNA-PK controls genotoxic stress-induced type I Interferon responses in absence of cGAS

Previous work has shown that in myeloid cells, DNA-PKcs does not induce genotoxic stress-associated type I IFN responses ^19^. Here, we questioned whether DNA-PK may be involved in genotoxic stress-associated type I IFN responses in cancer cells lacking cGAS. To this aim, T98G cells were treated with the camptothecin genotoxic agent to induce dsDNA breaks that are primarily repaired by NHEJ ^28^. Staining with a dsDNA-specific antibody showed that such treatment is sufficient to induce cytosolic accumulation of dsDNA (Figure 1H, upper panels) together with accumulation of 53BP1 foci in the nucleus and in the cytosol, reflecting defective repair and accumulation of DNA lesions (Figure 1H, lower panels) ^29^. WB analysis showed that camptothecin treatment led to increased pDNA-PKcs, pIRF3, but not pSTING (Figure 1I), together with increased levels of *IFNB* and *CXCL10* (Figure 1J). This suggests that DNA-PK may be responsible for cGAS/STING-independent type I IFN responses following genotoxic stress in T98G cells.

We next treated T98G cells with camptothecin in the presence of NU7441. Since NU7441 inhibits the catalytic activity of DNA-PKcs, as reflected by decreased pDNA-PKcs (Figure S1F), such treatment presumably impacts both DNA-PK-mediated DNA repair and signaling. Consequently, treatment with NU7441 alone was sufficient to promote the accumulation of DNA damage, as confirmed by the presence of 53BP1 foci and of cytosolic dsDNA (Figure S1G), without triggering type I IFN responses, as attested by absence of *IFNB* and *CXCL10* upregulation (Figure 1K). Treatment with NU7441 abrogated camptothecin-associated *IFNB* and *CXCL10* induction (Figure 1K), supporting that DNA-PKcs is responsible for genotoxic stress-dependent activation of type I IFN responses in T98G cells.

### DNA-PK-dependent detection of cytosolic dsDNA drives STING-independent IRF3-dependent type I IFN responses in cancer cells lacking cGAS

We next questioned the molecular mechanisms involved in DNA-PK-dependent type I IFN responses following challenge with exogenous dsDNA or genotoxic stress. First, immunofluorescence analysis showed that following dsDNA challenge DNA-PKcs and pDNA-PKcs are found in the cytosol (Figure 2A). Second, we assessed the ability of DNA-PK to interact with cytosolic dsDNA. To this aim, we used 80nt-long dsDNA or ssDNA, bearing a 5’ biotin on the sense strand to perform streptavidin-affinity pull-down experiments using whole cell extracts from T98G cells. We thereby observed that DNA-PKcs, KU80 and KU70 are recruited to dsDNA (Figure 2B). Similar experiments were performed following 6 h of biotinylated dsDNA transfection in the THP-1 human myeloid cell line (Figure S2A), a time point at which transfected dsDNA are found in the cytosol ^3^. In these conditions, together with DNA-PKcs, KU70 and KU80, recruitment of pDNA-PKcs to dsDNA was also observed (Figure S2A). Combined, these experiments support that DNA-PK is recruited to cytosolic dsDNA.

**Figure 2:**
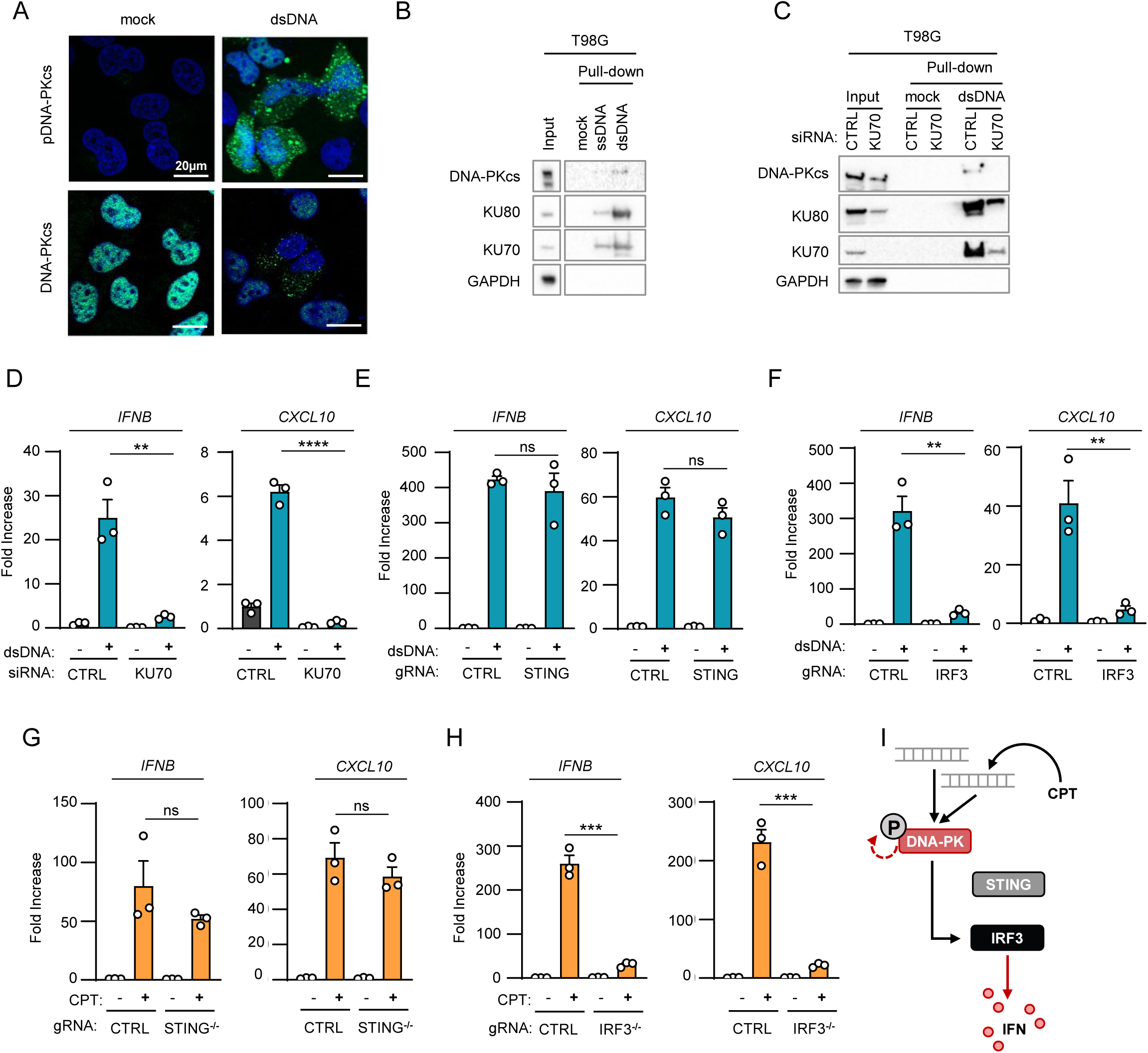
DNA-PK-dependent detection of cytosolic dsDNA drives STING-independent IRF3-dependent type I IFN responses in cancer cells lacking cGAS. (A) T98G cells were treated or not with dsDNA for 6 h prior to IF analysis using DNA- PKcs- and pDNA-PKcs-specific antibodies, and DAPI nuclear staining. Representative images are shown (n=3). (B) Whole cell extracts from T98G cells were incubated with 80nt-long biotinylated ssDNA or dsDNA prior to pull-down using streptavidin-affinity beads. Input and eluates were analyzed by WB using indicated antibodies. (C) T98G cells were treated with non-targeting (CTRL) or KU70-targeting siRNAs prior to whole cell extract preparation, and pull-down as in B. Inputs and eluates were analyzed by WB using indicated antibodies. (D) T98G cells were treated with CTRL or KU70-targeting siRNAs prior to challenge with dsDNA for 6 h. *IFNB* and *CXCL10* mRNA levels were analyzed by RT-qPCR. Graphs present a representative biological triplicate (n=3). (E) CTRL or *STING^-/-^* T98G cells were transfected or not with dsDNA for 8 h prior to analysis of *IFNB* and *CXCL10* mRNA levels (n=3). (F) As in E, except that CTRL or *IRF3*^-/-^ T98G cells were transfected or not with dsDNA for 6 h prior to analysis (n=3). (G) CTRL or *STING-/-* T98G cells were treated or not with CPT for 72 h prior to analyses of *IFNB* and *CXCL10* mRNA levels (n=3). (H) As in G, except that CTRL and *IRF3^-/-^* T98G cells were used. Graphs present a representative biological triplicate (n=3). (I) Schematic representation of cytosolic dsDNA-dependent type I IFN induction in T98G cells. All immunoblots are representative experiments. All graphs present means ± SEM. P- values were determined by Student’s t-test. ns: not significant. ** p < 0.01, *** p < 0.001, **** p < 0.0001.

We next tested whether interaction of KU70:KU80 with dsDNA was required for DNA-PKcs-dependent type I IFN responses. To this aim, we performed KU70 knock-down (Figure 2C and S2B), prior to assessment of the recruitment of DNA-PK to dsDNA. Knock-down of KU70 decreased the recruitment of KU80 while abolishing the recruitment of DNA-PKcs to dsDNA (Figure 2C). Conversely, performing dsDNA pull-down using whole cell extracts from control (HCT116^CTRL^) and DNA-PKcs-deficient HCT116 (HCT116*^PRKDC-/-^*) cells showed that DNA-PKcs is not required for the recruitment of KU70 and KU80 to dsDNA (Figure S2C). Importantly, knock-down of KU70 also abolished dsDNA-induced *IFNB* and *CXCL10* expression in T98G cells (Figure 2D). Thus, these data demonstrate that the recruitment of DNA-PKcs to cytosolic dsDNA through the KU70:KU80 heterodimer is required to trigger type I IFN responses.

WB analyses of STING protein levels and phosphorylation status in glioblastoma cells (Figure 1 and S1) suggest that the DNA-PK-dependent type I IFN responses elicited by dsDNA transfection is STING-independent. To formally test the requirement for STING, we assessed dsDNA-induced type I IFN responses in STING-deficient T98G cells (T98G*^STING-/-^*) (Figure 2E and S2D). Challenge with dsDNA of T98G*^STING-/-^*and of their control counterparts (T98G^CTRL^) promoted IRF3 phosphorylation (Figure S2D) together with *IFNB* and *CXCL10* upregulation (Figure 2E), regardless of STING expression. The presence of phosphorylated IRF3 following dsDNA challenge in cGAS-deficient cells (Figure 1A, E and S1B) also suggested that DNA-PK-associated IFN responses requires IRF3. To confirm this requirement, IRF3-deficient T98G cells (T98G*^IRF3-/-^*) (Figure S2E) were challenged with dsDNA prior to assessment of type I IFN responses. Absence of IRF3 disrupted type I IFN responses in T98G cells (Figure 2F and S2E), supporting that DNA-PK-dependent, STING-independent, type I IFN responses require IRF3 in cGAS-deficient cancer cells.

We next analyzed whether genotoxic stress-induced DNA-PK-dependent type I IFN responses in cGAS-deficient cells are governed by similar molecular mechanisms. To this aim, T98G^CTRL^, T98G*^STING-/-^* and T98G*^IRF3-/-^* were treated with camptothecin prior to analysis of DNA-PK activation and IFN responses. Similar to what was observed upon challenge with dsDNA, genotoxic stress induced DNA-PKcs phosphorylation, regardless of the expression of STING and IRF3 (Figure S2F-G). In addition, while induction of type I IFN responses did not require expression of STING (Figure 2G), the presence of IRF3 was required (Figure 2H). Thus, DNA-PKcs controls genotoxic stress-induced type I IFN responses through IRF3 activation.

Finally, we questioned whether cGAS-independent DNA-PK-associated type I IFN responses can be observed in other cell lines. To this aim, we used cGAS-knock-out THP-1 cells (THP-1*^cGAS-/-^*) and observed that, challenged with dsDNA, induced type I IFN responses (Figure S2H-I), concomitantly to DNA-PKcs phosphorylation (Figure S2H). However, STING ablation abolished type I IFN responses in THP-1 cells (Figure S2H-I). Thus, in contrast to T98G cells, STING is non-dispensable for type I IFN responses in myeloid cells, suggesting that the requirement for STING in DNA-PK activation is cell type-specific. Thus DNA-PKcs drives cytosolic dsDNA-IFN responses through IRF3 (Figure 2I).

### cGAS and DNA-PKcs cooperate for optimal dsDNA-induced type I Interferon responses

Considering that DNA-PK can elicit type I IFN responses in absence of cGAS, we next interrogated the impact of co-expressing DNA-PKcs and cGAS in glioblastoma cells. To this aim, we generated T98G cells stably expressing cGAS (T98G^cGAS^) or not (T98G^Empty^) (Figure 3A). Expression of cGAS was sufficient to induce constitutive degradation of STING (compare lanes 3 to 1) and to promote increased basal *IFNB* and *CXCL10* levels (Figure 3B). Additionally, challenge with dsDNA led to enhanced pIRF3 levels together with increased induction of type I IFN responses (Figure 3A-B), indicating that the cGAS-STING axis is efficiently restored in the T98G^cGAS^ cells. Knowing the strong affinity of cGAS for cytosolic dsDNA ^30^, we addressed whether DNA-PK and cGAS can compete for dsDNA detection, by transfecting biotinylated dsDNA in T98G^Empty^ and T98G^cGAS^ or in THP-1^CTRL^ and THP-1*^cGAS-/-^* prior to streptavidin-affinity pull-down. WB analysis showed that the recruitment of subunits of DNA-PK to dsDNA is not altered in the presence of cGAS (Figure 3C and S3A). Thus, expressing cGAS in T98G cells restores the cGAS-STING signaling axis without modifying the interaction of DNA-PK with dsDNA ligands.

**Figure 3:**
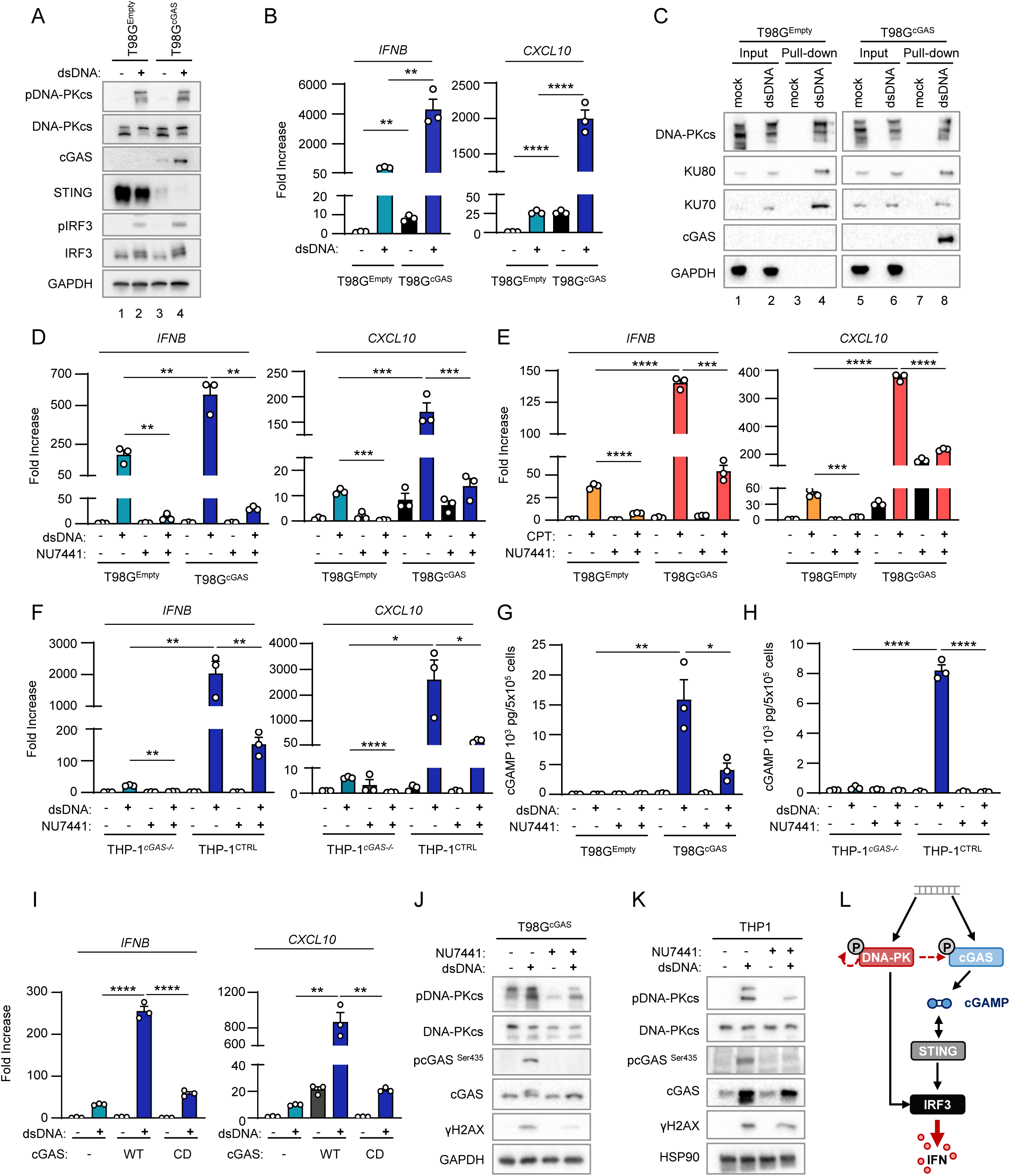
cGAS and DNA-PKcs cooperate for optimal dsDNA-induced type I Interferon responses. (A) T98G cells stably expressing cGAS (T98G^cGAS^) or not (T98G^Empty^) were transfected or not with dsDNA for 6 h prior to whole cell extraction and WB analysis using indicated antibodies. (B) *IFNB* and *CXCL10* mRNA levels were analyzed by RT-qPCR in samples treated as in A. Graphs present a representative biological triplicate (n=5). (C) T98G^Empty^ and T98G^cGAS^ were transfected or not with biotinylated dsDNA prior to whole cell extraction and pull-down using streptavidin-affinity beads. Inputs and eluates were analyzed by WB using indicated antibodies. (D) T98G^Empty^ and T98G^cGAS^ were transfected or not with dsDNA for 6 h in the presence or not of NU7441 prior to *IFNB* and *CXCL10* expression analysis. Graphs present a representative biological triplicate (n=5). (E) T98G^Empty^ and T98G^cGAS^ were treated or not with CPT for 72 h in combination or not with NU7441 (48 h) prior to *IFNB* and *CXCL10* expression analysis. Graphs represent a biological triplicate (n=4). (F) THP-1^CTRL^ and THP-1*^cGAS-/-^* were transfected or not with dsDNA for 6 h in presence or not of NU7441 prior to *IFNB* and *CXCL10* expression analysis (n=3). (G) Intracellular cGAMP levels were analyzed in samples treated as in E by ELISA (n=3). (H) Intracellular cGAMP levels were analyzed in samples treated as in F by ELISA. Graphs present a representative biological triplicate (n=2). (I) T98G expressing a catalytic dead cGAS allele (T98G^cGAS-CD^) and T98G were treated as in A prior to *IFNB* and *CXCL10* levels analysis. Graphs present a representative biological triplicate (n=3). (J) T98G^cGAS^ were transfected or not with dsDNA in presence or not of NU7441, prior to WB analysis using indicated antibodies. (K) THP-1 were transfected or not with dsDNA in presence or not of NU7441, prior to WB analysis using indicated antibodies. (L) Schematic representation of the molecular mechanisms involved in the cooperation between DNA-PKcs and cGAS for type I IFN induction. All immunoblots are representative experiments. All graphs present means ± SEM. P-values were determined by Student’s t-test. ns: not significant. * p < 0.05, ** p < 0.01, *** p < 0.001, **** p < 0.0001.

Given that both cGAS and DNA-PK can detect cytosolic DNA when co-expressed, we next questioned their respective contribution to dsDNA-dependent induction of type I IFN responses. To this aim, T98G^Empty^ and T98G^cGAS^ were either transfected with dsDNA or treated with camptothecin, in the presence or not of NU7441, prior to evaluation of type I IFN responses. Intriguingly, treatment with NU7441 led to a dramatic decrease of *IFNB* and *CXCL10* expression, both in the presence and absence of cGAS, following dsDNA transfection (Figure 3D and S3B). Similarly, treatment with camptothecin led to higher type I IFN responses in the presence of cGAS, that were abolished by treatment with NU7441 (Figure 3E). This suggested that DNA-PK and cGAS cooperate for the induction of type I IFN responses, when co-expressed. We interrogated whether a similar cooperation could be witnessed in other cell lines, using THP-1, but also the CFPAC pancreatic cancer cell line, and their cGAS^-/-^ counterparts. Treatment with NU7441 also led to a dramatic decrease of type I IFN responses to dsDNA in both cell types, regardless of the expression of cGAS (Figure 3F and S3C-E). Although we had observed that DNA-PK-dependent activation of dsDNA-related type I IFN responses requires STING in THP-1 cells, but not in T98G cells (Figure 2 and S2). Thus, our data suggest that the mechanism through which DNA-PK and cGAS synergize is common to both cell types.

To identify the molecular mechanism through which cGAS and DNA-PK cooperate, we tested whether DNA-PK can control cGAS activity. To this aim, we quantified intracellular cGAMP levels in T98G^Empty^ and T98G^cGAS^, but also in THP-1^CTRL^ and THP-1*^cGAS-/-^* cells, upon challenge with dsDNA in the presence or not of NU7441. Treatment with NU7441 led to a decrease of cGAMP levels in both T98G^cGAS^ and THP-1 cells (Figure 3G-H), although NU7441 did not alter cGAS activity *in vitro* (Figure S3F). Finally, when T98G stably expressing a catalytic-dead cGAS allele (T98G^cGAS-CD^) were challenged with dsDNA, type I IFN responses were drastically reduced as compared to those witnessed in T98G^cGAS^ (Figure 3I and S3G). These data show that the cooperation between cGAS and DNA-PK operates at the level of cGAS activity, and strongly suggest that DNA-PKcs boosts the production of cGAMP by cGAS.

Phosphorylation of the Serine 435 (Ser435) of cGAS has been previously shown to be required to enable cGAS-dependent cGAMP production ^31^. Because DNA-PKcs bears a Serine/Threonine kinase activity, we asked whether DNA-PKcs could be responsible for this phosphorylation. Challenge with dsDNA of cGAS-proficient cells (namely T98G*^cGAS^*, THP-1 and CFPAC) showed that the phosphorylation of cGAS on Ser435 was lost upon DNA-PKcs inhibition by NU7441 (Figure 3J, K and S3D). This supports that DNA-PKcs catalytic activity is required for cGAS phosphorylation at Ser435. Thus, altogether these data show that DNA-PKcs is required for efficient cGAS-dependent cGAMP production (Figure 3L).

### cGAS re-expression in glioblastoma cancer cells promotes macrophage recruitment and impairs tumorigenesis

We next questioned whether the synergy between cGAS and DNA-PK could have an impact on glioblastoma tumor immunogenicity. First, because cGAS has been previously shown to alter proliferation ^32, 33^, we assessed the growth of T98G^Empty^ and T98G^cGAS^ in 2D and 3D cultures. Follow-up over time showed that cGAS re-expression did not alter the proliferation of T98G cells (Figure 4A-B), nor affected their ability to form spheroids (Figure 4C), ruling out cell-intrinsic defects. However, upon subcutaneous engraftment in nude mice, T98G^cGAS^ failed to form tumors (Figure 4D-E). To visualize early steps of tumorigenesis, we next used zebrafish embryos in which we performed orthotopic transplantation of T98G^Empty^ and T98G^cGAS^ stably expressing a *green fluorescence protein* (GFP) reporter (T98G-GFP^Empty^ and T98G-GFP^cGAS^). Monitoring of the intracranial GFP signal over time showed a faster decrease of the T98G-GFP^cGAS^ tumor mass, as compared to T98G-GFP^Empty^ tumors (Figure 4F). Moreover, morphological assessment of tumors showed more elongated pseudopodia, which are hallmarks of invasiveness ^34^, in T98G-GFP^Empty^ tumors as compared to T98G-GFP^cGAS^ (Figure S4A-B). Thus, cGAS expression in T98G cells is sufficient to impair early tumorigenesis.

**Figure 4:**
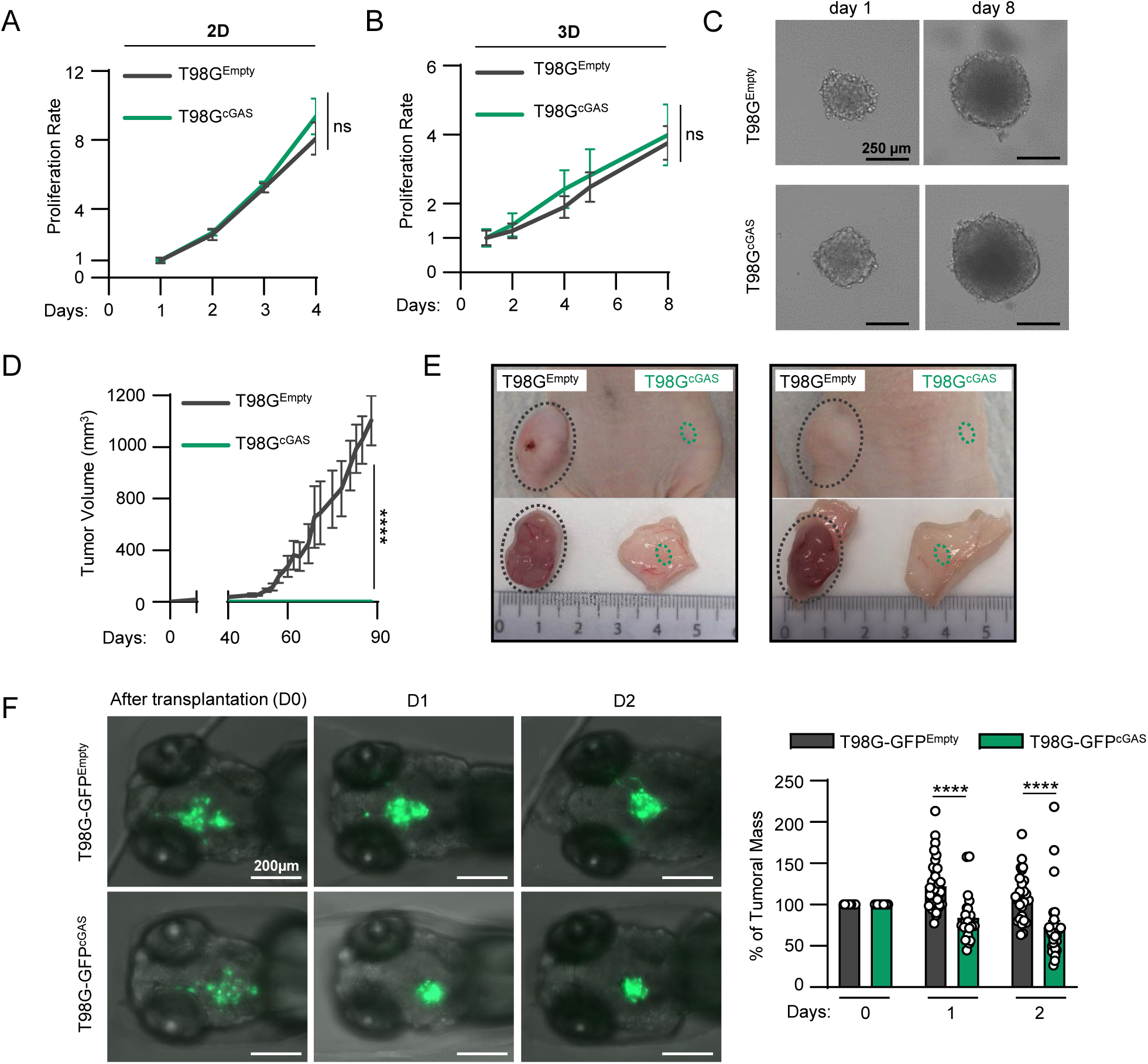
cGAS re-expression in glioblastoma cancer cells impairs tumorigenesis. (A) The proliferation of T98G^Empty^ and T98G^cGAS^ was monitored in 2D cultures over 4 days (n=3). (B) The volume of spheroids formed by T98G^Empty^ and T98G^cGAS^ was monitored over 8 days (n=3). (C) Representative images of T98G^Empty^ and T98G^cGAS^ spheroids at day 1 and day 8. (D) The volume of subcutaneous T98G^Empty^ and T98G^cGAS^ tumors in nude mice was measure every 3-4 days by caliper (n=6 mice per group). (E) Representative pictures of T98G^Empty^ and T98G^cGAS^ tumors at day 90 post subcutaneous engraftment. (F) T98G^Empty^ and T98G^cGAS^ stably expressing a GFP reporter (T98G-GFP^Empty^ and T98G-GFP^cGAS^, respectively) were xenotransplanted into the head of *tg(mfap4:RFP)* zebrafish line at 3 days post fertilization (dpf). Zebrafish embryos were imaged daily over 3 days. The graph represents the mean (±SEM) percentage of tumor growth normalized by the area on the day of transplantation (n=21 T98G-GFP^Empty^ and n=29 T98G-GFP^cGAS^ embryos). All graphs present means ± SEM. P-values were determined by Student’s t-test. ns: not significant. * p < 0.05, ** p < 0.01, **** p < 0.0001. One-way Anova with Tukey’s multiple comparisons test was used for mice analyses. Mann-Withney test was performed to analyze tumor growth in zebrafish.

Since nude mice and zebrafish embryos, at the stage at which they were engrafted, do not possess an adaptive immune system, we next hypothesized that differential cytokine and chemokine production may modulate myeloid cell activity in the glioblastoma microenvironment. Migration assays showed that conditioned media from T98G^cGAS^ cells increased the migration of THP-1 cells as compared to that from T98G^Empty^ cells (Figure 5A). However, conditioned media from T98G^cGAS^ cells was not sufficient to promote THP-1 and primary myeloid cell polarization (Figure S5A-B). Thus, soluble factors secreted by T98G^cGAS^ are sufficient to promote the recruitment of myeloid cells to the tumor mass, but not their polarization. To identify those soluble factors, we profiled cytokines and chemokines levels in the supernatant of T98G^Empty^ and T98G^cGAS^, thereby identifying C-C Motif Chemokine Ligand 2 (CCL2) and 5 (CCL5), in addition to CXCL10, as most upregulated in the supernatant of T98G^cGAS^ as compared to T98G^Empty^ cells (Figure 5B). Such upregulation was also observed at the gene expression level (Figure 5C) and was lost when comparing T98G^cGAS^ to T98G^cGAS^*^-^*^CD^ (Figure S5C). Thus, these data support that the synergy between DNA-PK and cGAS promotes the production of chemokines that can induce macrophage recruitment.

**Figure 5:**
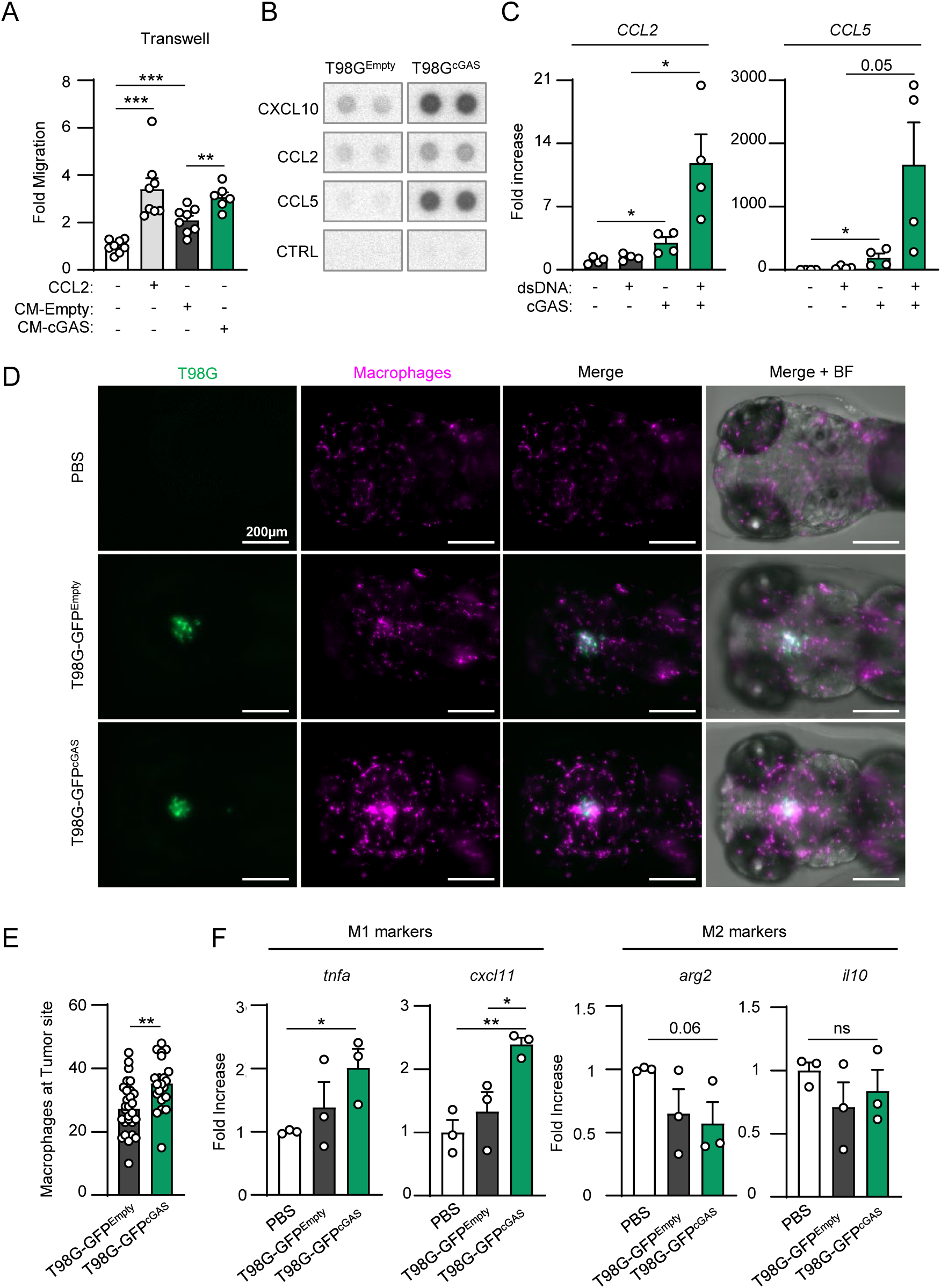
cGAS re-expression in glioblastoma cancer cells promotes macrophage recruitment. (A) Graph represents the mean (± SEM) fold migration of THP-1 cells through a 3 µm transwell insert when conditioned media from T98G^Empty^ or T98G^cGAS^ was applied to lower chamber for 6 h. CCL2 was used as positive control (n=8). (B) CXCL10, CCL2 and CCL5 protein levels in conditioned media from T98G^Empty^ and T98G^cGAS^ cells were assessed using proteome profiler. Proteins that were found to be the most upregulated in T98G^cGAS^ are shown. Representative immunoblots. (C) T98G^Empty^ and T98G^cGAS^ were transfected or not with dsDNA prior to analyses of *CCL2* and *CCL5* by RT-qPCR (n=4). (D) Zebrafish embryos injected with T98G-GFP^Empty^, T98G-GFP^cGAS^, or PBS at 3 dpf (Figure 4E) were imaged at 24 h post transplantation. Representative images of macrophage recruitment (purple) in the head (n=21 T98G-GFP^Empty^ and n=29 T98G-GFP^cGAS^ embryos). (E) Graph presents the quantification of macrophages recruited at tumor site 24 h post xenotransplantation in D. (F) Heads of zebrafish treated as in D were isolated prior to RNA extraction and analysis of M1 (*tnfa* and *cxcl11*) or M2 (*arg2* and *il10*) polarization markers. Each value in the graph is the mean of 25 embryos. All graphs present means ± SEM. P-values were determined by Student’s t-test. ns: not significant. * p < 0.05, ** p < 0.01, **** p < 0.0001. Mann-Withney test was performed to analyze macrophage recruitment in zebrafish.

We thus took advantage of the optical traceability of macrophages in the *tg(mfap4:RFP)* zebrafish line, owing to the expression of a red fluorescent protein (RFP) reporter under the control of the *mfap4* promoter. Imaging over time showed enhanced recruitment of myeloid cells around and inside tumors of zebrafish embryos injected with T98G-GFP^cGAS^, as compared to those injected with T98G-GFP^Empty^ (Figure 5D). Quantification of contacts between tumors and myeloid cells further supported increased recruitment and interaction between tumor masses formed of T98G-GFP^cGAS^ and myeloid cells, as compared to T98G-GFP^Empty^ tumor masses (Figure 5D-E). In addition, assessment of macrophage polarization markers showed increased expression of M1 markers in zebrafish heads where T98G-GFP^cGAS^ were injected, as compared to T98G-GFP^Empty^ tumors (Figure 5F). Thus, cGAS re-expression in T98G cells is sufficient to promote macrophage recruitment and M1 polarization at the tumor site, impairing tumor engraftment and promoting tumor clearance.

### cGAS and DNA-PKcs levels increase with tumor grade

To confirm the physiological relevance of our findings that indicate that cGAS expression impairs early tumorigenesis, we performed meta-analysis of glioblastoma tumors, using the GlioVis database ^35^. We first examined the co-expression of *PRKDC* and *MB21D1* (encoding cGAS) in glioblastoma tissue samples from the CGGA transcriptomic database. Of the 224 retrieved cases, 3 populations were determined based on *PRKDC* and *MB21D1* expression levels: *PRKDC*^low^/*MB21D1*^low^, *PRKDC*^high^/*MB21D1*^low^ and *PRKDC*^high^/*MB21D1*^high^ (Figure 6A). Analysis of mRNA levels of total macrophages, chemokines/cytokines and macrophage polarization markers were performed between these groups. We found that expression of *MB21D1* correlated with high expression of *CXCL10*, *CCL2* and *CCL5* chemokines (Figure 6B) and with the increased presence of macrophages (Figure 6C-E). Consistent with Pan-cancer analysis ^15^, analysis of patient survival showed that higher *MB21D1* expression led to worst patient survival as compared to patients with low *MB21D1* expression, supporting that expression of cGAS is a poor outcome marker in glioblastoma (Figure 6F). In addition, analysis of *PRKDC* and *MB21D1* expression in glioblastoma of grades II, III and IV, indicates that *PRKDC* and *MB21D1* expression increased significantly with the aggressiveness of the tumors (Figure 6G). Thus, altogether, our data support that DNA-PK and cGAS cooperate to foster a pro-inflammatory environment, through enhancing the production of cytokines and chemokines that attract macrophages to the tumor vicinity, a process that inhibits early tumorigenesis, but fuels cancer-associated inflammation at later stages.

**Figure 6:**
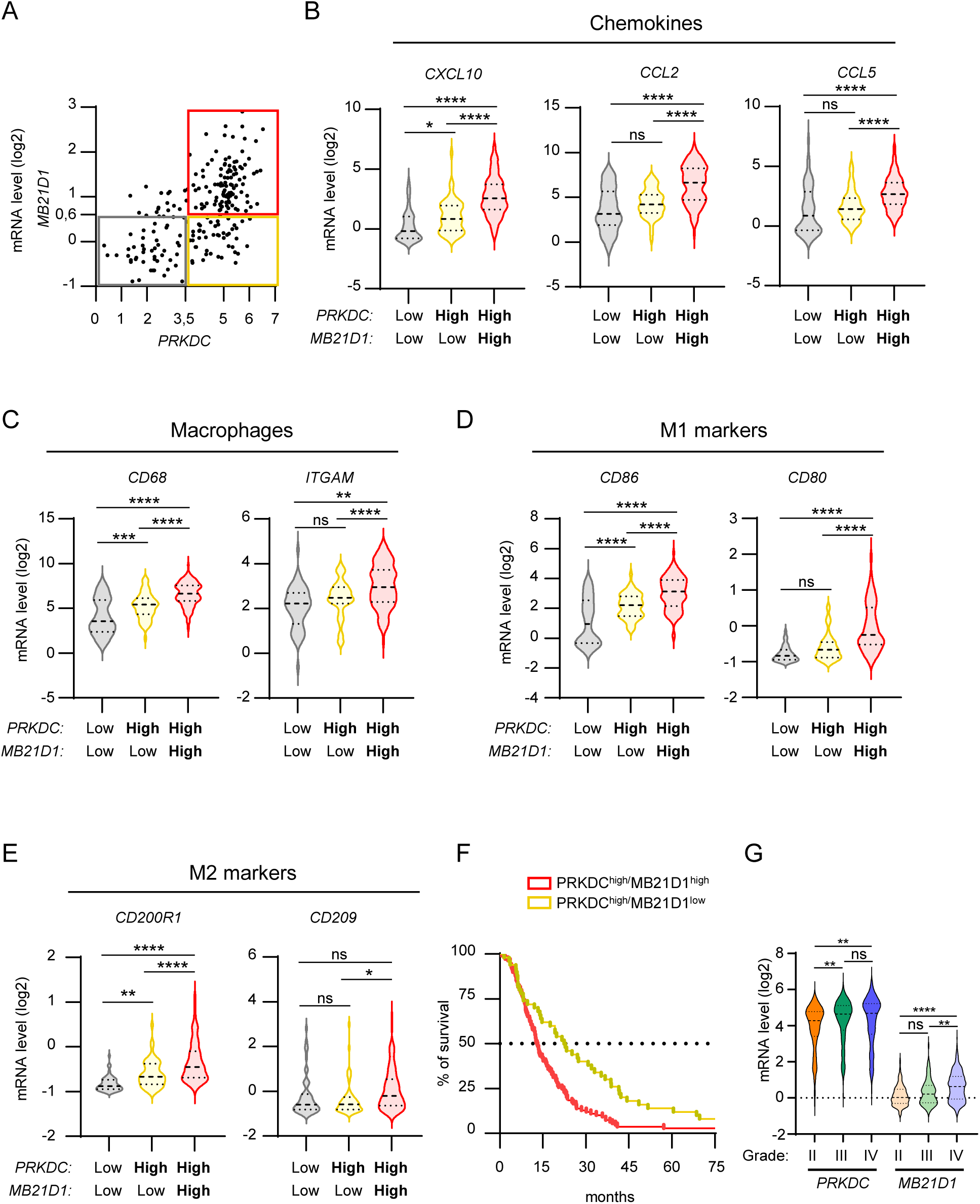
cGAS and DNA-PKcs levels increase with tumor grade. (A) Correlation plot between *PRKDC* and *MD21B1* expression in glioblastoma patients. Three distinct populations can be visualized: *PRKDC^low^*/*MD21B1^low^*; *PRKDC^high^/MD21B1^low^* and *PRKDC^high^/MD21B1^high^* (total patients n=224). (B) Violin plots present chemokine gene expression (*CXCL10*, *CCL2* and *CCL5*) in glioblastoma samples from A. (C) Violin plots present macrophage gene expression (*CD68* and *ITGAM*) in glioblastoma samples from A. (D) Violin plots present pro-inflammatory M1 macrophage gene expression (*CD86* and *CD80*) in glioblastoma samples from A. (E) Violin plots present anti-inflammatory M2 macrophages gene expression (*CD200R1* and *CD209*) in glioblastoma in samples from A. (F) Graph presents the survival rate of glioblastoma patients from A that present *PRKDC^high^/MB21D1^high^* and *PRKDC^high^/MB21D1^low^* expression. (G) Violin plots present the expression of *PRKDC* and *MB21D1* in datasets analyzed in A, based on tumor grade (II to IV). All graphs present means ± SEM. P-values were determined by Student’s t-test. ns: not significant. * p < 0.05, ** p < 0.01, *** p < 0.001, **** p < 0.0001. One-way Anova with Tukey’s multiple comparisons test was used to compare gene expression among populations in glioblastoma dataset.

## DISCUSSION

We demonstrate that DNA-PK and cGAS synergize for the production of type I IFNs and chemokines, thus dictating the composition of the tumor microenvironment. This cooperation is therefore an attractive target to modulate tumor immunogenicity. Our data further reveal that the molecular determinants of the activation of DNA-PK-dependent signaling is governed by cell-type specific rules, that, if adequately harnessed, may allow targeting inflammatory responses in specific cells of the tumor microenvironment.

Tumor-associated macrophages are the most abundant immune cell population in the glioblastoma tumor microenvironment ^36^ and their presence is generally an indicator of poor outcome for glioblastoma patients ^37^. In agreement, blocking macrophage recruitment through *Ccl2* genetic ablation ameliorates mice survival ^38^. Moreover, patients with low tumoral *CCL2* expression survived significantly longer than those with high *CCL2*^38^. Since pDNA-PKcs expression positively correlates with tumor progression ^39^, and in the light of our meta-analysis, it is tempting to hypothesize that in tumors where the cGAS-STING pathway fuels tumorigenesis, the use of DNA-PKcs inhibitor may facilitate tumor clearance.

Indeed, DNA-PK inhibitors have been used in preclinical studies in glioblastoma, bringing promising results ^39, 40^ and several clinical trials are ongoing (NCT02977780 and NCT04555577). To fully exploit the benefit of DNA-PKcs inhibition, our data support that the expression of cGAS is an important parameter to consider. Conversely, DNA-PK agonists could allow the re-establishment of inflammatory responses in tumors in which the cGAS pathway is not functional, or boost anti-tumoral inflammatory responses in those expressing cGAS. This approach may represent a promising therapeutic avenue in glioblastoma patients where STING agonists have shown benefits ^41^. Yet, the use of STING agonists has shown cell-type specific drawbacks that should not be overlooked ^42–44^. In the same line, our data support that exploring the functionality of DNA-PK and the impact of STING activation in the different cell types composing the tumor microenvironment is critical for patient stratification.

Using transgenic zebrafish line, we revealed that the presence of cGAS in tumor cells, at early stages, is sufficient to elicit myeloid cell recruitment and polarization into M1 macrophages that are key players in the initiation of antitumor responses. However, high levels of cGAS and STING predict poor prognosis ^15^, as supported by our glioblastoma patient data meta-analysis. This observation, together with the fact that terminally differentiated healthy tissues, do not express a functional cGAS-STING signaling axis ^45^, supports that cGAS expression is acquired during tumorigenesis, to the contrary of prior assumptions that cGAS downregulation may be an immune escape mechanism.

Our study raises the possibility that in inflammatory pathologies presenting with pathological chronic STING activation, inhibition of DNA-PKcs in combination with classical Janus kinase inhibitors, that are already used in standard patient care ^46^, may allow better suppression of chronic type I IFN responses. Conversely, *PRKDC* mutations are associated with auto-inflammatory pathologies ^47, 48^ in which cGAS-STING activation should be explored for the design of novel therapeutic avenues.

## ACKNOWLEDGMENTS

We thank P. Pourquier for the HCT116 cell lines, N. Bonnefoy for the parental CFPAC cell line, Soren Paludan for THP-1 cell lines. We thank C. Goujon for the parental T98G cell line and CRISPR/Cas9 gRNA sequences. We thank C. Langevin and all members of the molecular basis of inflammation laboratory for discussions. We thank Noemie Dupuis and Aischa Blömeke-Eiben for technical assistance.

We acknowledge iExplore-RAM animal facility, the MRI imaging facility, member of the national infrastructure France-BioImaging infrastructure supported by the French National Research Agency (ANR-10-INBS-04, “Investments for the future”) and the SIRIC Montpellier Cancer (INCa_Inserm_DGOS_12553). Work in NL’s laboratory was funded by the European Research Council (ERC-Stg CrIC: 637763, ERC-PoC DIM-CrIC: 893772), LA LIGUE pour la recherche contre le cancer, Agence Nationale de Recherche sur le SIDA et les Hépatites virales (ANRS: ECTZ117448), La Région Languedoc Roussillon, and the CNRS. CT was supported by the Merck Sharp and Dohme Avenir (MSD-Avenir – GnoSTic) program and ANRS (ECTZ119088). HC was supported by LA LIGUE pour la recherche contre le cancer. SG was supported by a LabMuse EpiGenMed. ALCV was supported by ERC-Stg CrIC: 637763, and MSD-Avenir. KP was supported by the Labex EpiGenMed. IKV was supported by ERC-Stg CrIC: 637763, followed by Fondation pour la Recherche Médicale (FRM: ARF20170938586 “Prix Roger PROPICE pour la recherche sur le cancer du pancreas”. JM was supported by an Agence Nationale de Recherche Technologie” (ANRT).

## AUTHOR CONTRIBUTIONS

Conceptualization: NL, CT. Methodology: NL, CT, IKV, JM, FB, DP, KP ALCV. Investigation: CT, JM, IKV, KP, ALCV. Visualization: NL, CT, JM, IKV. Supervision: NL, JPH, FB, KK, LF. Writing—original draft: NL, CT, JM. Writing—review & editing: NL, CT, JM, IKV.

## MATERIALS AND METHODS

### Resource availability

Requests for resources and reagents should be directed to Dr. Nadine Laguette.

### Animals

Experiments in mice were conducted using Athymic Nude Foxn1-nu males in which subcutaneous injections were performed, prior to follow-up of tumor size over time using a caliper. These experiments were performed in agreement with European rules and regulations for animal handling (25066-2020040315236430).

*In vivo* experiments in zebrafish were performed using the *tg(mfap4:RFP)* zebrafish line. Each experiment was conducted using at least 21 individual fish per condition, as indicated in the figure legend. All experimental procedures on zebrafish were performed in accordance with the European guidelines and regulations for Animal Protection and authorization no. F341725 from the French Ministry of Health.

### Cells and cell cultures

THP-1^CTRL^, THP-1*^cGas−/−^* and THP-1*^STING−/−^* were a gift of S. R. Paludan, HCT116^CTRL^ and HCT116*^PRKDC-/-^* were obtained from P. Pourquier, while parental T98G and CFPAC were provided by from C. Goujon and N. Bonnefoy, respectively. Gli4 and Gli7 were a gift from J.-P. Hugnot.

293T, T98G and their genetically engineered derivatives, and CFPAC and their genetically engineered derivatives, were maintained in Dulbecco’s modified Eagle’s medium (DMEM) supplemented with 10% fetal bovine serum (FBS, Eurobio), 1% penicillin/streptomycin (Lonza), and 1% L-glutamine (Lonza). HCT116, THP-1 cells and their derivatives were cultured in RPMI media (Lonza) supplemented with 10% FBS, 1% penicillin/streptomycin and 1% L-glutamine.

Human glioblastoma cancer stem cells Gli4 and Gli7 were cultured in T75 tissue culture flasks precoated with 40 μg/cm^2^ of poly-2-hydroxyethyl methacrylate (poly-HEME, Sigma) to avoid cell adhesion. They were cultivated in DMEM/F12 1:1 (Invitrogen), N2 and B27 supplements (Invitrogen), 2 mM glutamine (Invitrogen), 0.6% glucose (Sigma), 20 μg/mL bovine insulin (Sigma) supplemented by 2 μg/mL Heparin (Sigma), 20 ng/mL EGF (Peprotech) and 10 ng/mL FGF2 (Peprotech).

### Compounds

Nu7441: PubChem SID 249565690

Camptothecin: PubChem CID 24360

### Viral particle production and transduction

To generate knockout cell lines and the corresponding control cell lines, lentiviral particles were produced by co-transfection of 2 × 10^6^ 293T cells with 5 μg of LentiCRISPRv2GFP or LentiCRISPR v2 plasmids expressing the gRNA targeting the gene of interest or non-targeting control (CTRL) gRNA, 5 μg of psPAX2 and 1 μg of pMD2.G, using the standard calcium phosphate transfection protocol.

To generate the T98G cell lines stably expressing WT-cGAS, cGAS-CD or the corresponding control cell line, retroviral particles containing the transgene encoding Flag-and HA-tagged cGAS (F/HA-cGAS) alleles were produced by co-transfecting 1 × 10^6^ 293T cells with 5 μg of pOZ-F/HAcGAS or pOZ-F/HAcGAS-CD or pOZ-F/HA, 2.5 μg of MLV GagPol, and 2.5 μg of A-MLV envelope. Viral particles were harvested 48 h after transfection, filtered with 0.45 μM filters, and used for transduction.

For the generation of stable cell lines, 6 × 10^5^ T98G cells were seeded 24 h prior to transduction. Medium was replaced 8 h post transduction. Selection was performed 72 h post transduction using 2 µg/mL puromycin for at least 3 days. Selected cells were amplified and the levels of the protein of interest analyzed by western blot (WB).

### Generation of knock-out cell lines

T98G*^IRF3-/-^* and T98G^CTRL^ cell lines were generated using the LentiCRISPRv2GFP plasmid (gift from David Feldser; Addgene # 82416). IRF3 or control guide RNAs were inserted into the LentiCRISPRv2GFP plasmid following provided guidelines and lentiviral particles generated. T98G cells were transduced with lentiviral particles and 72 h post-transduction GFP positive cells were sorted and pooled in a 6 well plate using a BD FACS melody. Cells were next amplified and levels of IRF3 controlled by WB.

Generation of T98G*^STING-/-^*and T98G^CTRL^ cell lines was conducted as above, except that the LentiCRISPR v2 plasmid (Addgene #5296) was used and cells were selected 72 h post transduction using 2 µg/mL puromycin for 7 days. Cells were subsequently amplified and expression of STING controlled by WB. CFPAC*^cGAS-/-^* and CFPAC^CTRL^ cell lines were generated using a similar protocol except that puromycin-selected cells were further subjected to clonal selection using limiting dilutions. Clones were subsequently selected based on cGAS protein levels as evaluated by WB.

Guide RNAs used in this study:

For the generation of the IRF3 knockout cell line:

CTRL fwd: CACCGAGCACGTAATGTCCGTGGAT

CTRL rev: AAACATCCACGGACATTACGTGCTC

IRF3 fwd: CACCGAGCTGACACTCACCTTCCCC

IRF3 rev: AAACGGGGAAGGTGAGTGTCAGCTC

For the generation of the STING and cGAS knockout cell lines:

CTRL fwd: CACCGACGGAGGCTAAGCGTCGCAA

CTRL rev: AAACTTGCGACGCTTAGCCTCCGTC

STING fwd: CACCGCATATTACATCGGATATCTG

STING rev: AAACCAGATATCCGATGTAATATGC

cGAS fwd: CACCGGAACTTTCCCGCCTTAGGCA

cGAS rev: AAACTGCCTAAGGCGGGAAAGTTCC

### Generation of cell lines stably expressing cGAS

T98G overexpressing F/HA-cGAS (T98G^cGAS^) or a catalytic dead cGAS allele (T98G^cGAS- CD^), and their control cell line (T98G^Empty^) were generated by transducing parental T98G with retroviral particles produced by using the pOZ-F/HAcGAS; pOZ-F/HAcGAS-CD construct or empty vector, respectively, and selected with 2 µg/mL puromycin for 7 days.

### Generation of fluorescent glioblastoma cell line for zebrafish experiments

To obtain T98G-GFP^Empty^ and T98G-GFP^cGAS^, T98G^Empty^ and T98G^cGAS^ cell lines were stably transfected with sfGFP-N1 (gift from Michael Davidson & Geoffrey Waldo; Addgene #54737) using phosphate calcium. After transfection, cells were selected using Geneticin (800 µg/mL) for 4-6 weeks prior to zebrafish experiments.

### Site directed mutagenesis

To generate the catalytic dead mutant of cGAS, site directed mutagenesis was performed using the Quickchange Lightning kit of Agilent technologies, following the manufacturer’s instruction. Primers used for the mutagenesis reaction (Fwd: 5’-ggcggttttcacgtgatagtcgctgaacttgtccaagtgt-3’; rev: 5’-acacttggacaagttcagcgactatcacgtgaaaaccgcc-3), designed using the QuickChange Primer Design tool available online at https://www.agilent.com/store/primerDesignProgram.jsp, were purchased from Eurofins Genomics.

### Gene silencing

Silencing of KU70 and PRKDC was achieved in T98G cells using siRNAs and INTERFERin (Polyplus) following the manufacturer’s instructions.

The following siRNA sequences obtained by Dharmacon™ - Horizon Discovery were used:

si*CTRL*: CGUACGCGGAAUACUUCGAUU

si*PRKDC*: GAUCGCACCUUACUCUGUUUU

si*KU70*: ACAAGCAGUGGACCUGACUU

### Oligo sequences for annealing

To generate non-biotinylated or biotinylated dsDNA probes, dsDNA annealing was performed as described in ^3^. The following single strand probes obtained by IDT were annealed ^49^:

Fwd: ACATCTAGTACATGTCTAGTCAGTATCTAGTGATTATCTAGACATACATGATCTATGACATATATAGTGGATAAGTGTGG

Rev: CCACACTTATCCACTATATATGTCATAGATCATGTATGTCTAGATAATCACTAGATACTGACTAGACATGTACTAGATGT

### Whole-cell extract preparation and immunoblot

Cells were lysed in 5 packed cell volume of TENTG-150 [20 mM tris-HCl (pH 7.4), 0.5 mM EDTA, 150 mM NaCl, 10 mM KCl, 0.5% Triton X-100, 1.5 mM MgCl_2_, and 10% glycerol, supplemented with 10 mM β-mercaptoethanol and 0.5 mM PMSF] for 30 min at 4°C. Lysates were centrifuged 30 min at 13,000 rpm, and supernatants were collected for WB. For phosphorylated protein analysis, buffer was supplemented with PhosphoSTOP (Sigma) before whole-cell extraction. Protein quantification was performed using Bradford assay (Bio-Rad). Samples were run on either 4–15% Mini-PROTEAN® TGX™ Precast Protein Gels (Bio-Rad) (when analysis of DNA-PKcs and γH2AX was required) or on NuPAGE 10% or 12%, Bis-Tris Mini Protein gels (Invitrogen). Proteins were transferred onto nitrocellulose membranes. Membranes were incubated with primary antibodies (1:1000 dilution except when indicated) for 2 h at RT or over-night at 4°C. Primary antibodies used include: anti-pDNA-PKcs Ser2056 (ab124918, Abcam), anti-DNA-PKcs (A300-517AT, Bethyl, 1:500), anti-KU70 (4104S, Cell Signaling Technology), anti-KU80 (2753S, Cell Signaling Technology), anti-pcGAS Ser420 (AP1228, Abclonal), anti-cGAS (15102, Cell Signaling Technology), anti-pSTING Ser366 (191781, Cell Signaling Technology), anti-STING (13647, Cell Signaling Technology), anti-pIRF3 Ser386 (ab76493, Abcam), anti-IRF3 (11904, Cell Signaling Technology), anti-γH2AX (9718, Cell Signaling Technology), anti-HSP90 (4877, Cell Signaling Technology), anti-GAPDH (60004-1-Ig, Proteintech Europe, 1:5000), anti-αTUBULIN (66031-1-Ig, Proteintech Europe, 1:10000). Membranes were incubated with secondary antibodies (Cell Signaling Technology) at 1:2000 dilution, for 1 h at RT. Signal was visualized with SuperSignal West Pico Chemiluminescent Substrate (Thermo Fisher Scientific) or SuperSignal West Femto Maximum Sensitivity Substrate (Thermo Fisher Scientific), and images were acquired on a ChemiDoc (Bio-Rad) or using Amersham Hyperfilm™ ECL (GE Healthcare) films.

### Biotinylated nucleic acid pull-down using cell extracts following dsDNA transfection

Interaction of endogenous proteins and transfected biotinylated nucleic acids was assessed by transfecting T98G and THP-1 cells with nucleic acids (1 μg/ml) using JetPrime, according to the manufacturer’s protocol. Six hours after transfection, cells were harvested and lysed in TENTG-150 on ice for 30 min. Lysates were centrifuged at 13,000 rpm for 30 min at 4°C. Equal amounts of whole-cell lysates were incubated for 3 hours at 4°C on a wheel with 30 μl Dynabeads M280 pre-blocked in [100 mM NaCl, 2 mM DTT, 20 mg/mL BSA] overnight at 4°C on a wheel. After three washes in buffer [20 mM tris-HCl (pH 7.4), 0.5 mM EDTA, 0.05% Triton, 0.1% Tween, 150 mM NaCl, 10% glycerol, and 5 mM MgCl_2_], bound material was eluted in 30 µl Laemmli buffer. Protein interaction with the transfected biotinylated nucleic acids was assessed by WB.

### *In vitro* biotinylated nucleic acid pull-down

Pull-down was carried out using 30 μl of Dynabeads M280 per condition. Beads were blocked overnight as described above. After three washes in Washing Buffer [5 mM Tris-HCl (pH 7.5), 1 mM EDTA, 2 M NaCl], 3 µg of nucleic acids was coupled to 30 µl of beads according to the manufacturer’s instructions before equilibration in TENTG-150. Beads were then washed once with Washing buffer and equilibrated in TENTG-150. Whole cell extracts were diluted in TENTG-150. One milliliter of diluted lysate was added to the beads and incubated at 4°C on a wheel for 3 h in low-binding tubes (Axygen). Three consecutive washes were performed as above. Bound material was eluted in 30 μl of Laemmli buffer. Protein interaction with the biotinylated nucleic acids was assessed by WB.

### RNA extraction and RT-qPCR

RNA was extracted using TRIzol (Invitrogen) and treated with TURBO DNase (Ambion) according to manufacturer’s protocols. RNA was quantified with a Nanodrop spectrophotometer (ND-1000, Nanodrop Technologies). RNA (1-2 µg) was reverse transcribed using SuperScript IV reverse transcriptase (Invitrogen). Expression of specific mRNAs was determined with a LightCycler 480 (Roche) using the SYBR green PCR master mix (Takara). Reactions were performed in duplicate or triplicate, and relative amounts of cDNA were normalized to Glyceraldehyde3-phosphate dehydrogenase (*GAPDH*) for human cells or eukaryotic translation elongation factor 1 alpha 1, like 1 (*ef1a*) for zebrafish analyses.

Primers used for RT-qPCR analysis: Human:

*GAPDH* fwd: CTGGCGTCTTCACCACCATGG

*GAPDH* rev: CATCACGCCACAGTTTCCCGG

*IFNB* fwd: GAATGGGAGGCTTGAATACTGCCT

*IFNB* rev: TAGCAAAGATGTTCTGGAGCATCTC

*CXCL10* fwd: GAAAGCAGTTAGCAAGGAAAGGTG

*CXCL10* rev: ATGTAGGGAAGTGATGGGAGAGG

*CCL2* fwd: AGAATCACCAGCAGCAAGTGTCC

*CCL2* rev: TCCTGAACCCACTTCTGCTTGG

*CCL5* fwd: CCTGCTGCTTTGCCTACATTGC

*CCL5* rev: ACACACTTGGCGGTTCTTTCGG

Zebrafish:

*ef1a fwd:* AGAAGGCTGCCAAGACCAAG

*ef1a rev:* AGAGGTTGGGAAGAACACGC

*tnfa* fwd: GGAGAGTTGCCTTTACCGCT

*tnfa* rev: CCTGGGTCTTATGGAGCGTG

*cxcl11* fwd: ACTCAACATGGTGAAGCCAGTGCT

*cxcl11* rev: CTTCAGCGTGGCTATGACTTCCAT

*arg2* fwd: GAAGCCGTTCCTGTCTGCCA

*arg2* rev: TCGGCCTTTGCTTCCTTGCC

*il10* fwd: TCAGAGCAGGAGAGTCGAATGCA

*il10* rev: CGATTGGGGTTGTGGAGTGCTT

### cGAMP ELISA

For cGAMP quantification, T98G^Empty^, T98G^cGAS^, THP-1^CTRL^ and THP-1*^cGAS-/-^* were seeded 18 h before dsDNA transfection. One hour before dsDNA transfection cells were pre-treated with 2 µM NU7441 (#3712, Biotechne/Tocris) in OptiMEM and were subsequently transfected with dsDNA (1 µg/ml). Cells were harvested 6 h post transfection, counted, washed in phosphate-buffered saline (PBS) (Sigma), pelleted, and frozen at −80°C until extraction. cGAMP extraction was performed using the commercially available Mammalian Protein Extraction Reagent (M-PER) buffer (Thermo Fisher), accordingly to the manufacture protocol. The recovered supernatants were used for cGAMP measurement, following adequate sample dilution. cGAMP enzyme-linked immunosorbent assay (ELISA) was performed according to the manufacturer’s protocol using the Cayman Chemical 2′3′-cGAMP ELISA Kit (Bertin Bioreagents).

### Cell treatment and transfection

dsDNA transfections were conducted using previously published protocols ^50^. In brief, cells were plated in 6 well plate, B10 or B15 18 h before transfection. The day of transfection, media was carefully removed, plates were washed once with 1X PBS (room temperature) and 2, 10 or 20 ml Opti-MEM added, depending on the plate size. 2, 10 or 20 µg of dsDNA was transfected with the JetPrime transfection reagent (Polyplus) at 1:2 ratio. 6 h after transfection, cells were harvested and stored at -80°C prior to protein or RNA extraction.

To perform DNA-PKcs inhibition followed by dsDNA transfection, cells were pretreated with 2 µM of NU7441 in Opti-MEM, one hour prior the transfection.

When transfection was performed on Gli4 or Gli7, glioblastoma spheres were dissociated in Trypsin (0.2%, Sigma) at 37°C for 4 min. Trypsin inhibitor (Sigma, 50mg/mL), DNase I (0.015%, Roche) and CaCl_2_ (20mM) were subsequently added. After mechanical dissociation, cells were resuspended in PBS 1X. Following 2 washes with PBS 1X, cells were counted and 1 × 10^6^ cells were plated per well of a 6 well plate, precoated with poly-D-lysine (25 μg/mL) and laminin (2 μg/cm^2^, Sigma).

For chemotherapy treatment, cells were treated with 0.16 µM camptothecin (CPT) in DMEM for 48 or 72 h. When the treatment was in combination with the DNA-PKcs inhibitor, 2 µM of NU7441 was added 24 h and 48 h post CPT treatment.

### Conditioned media

For conditioned media preparation, 3.5 × 10^6^ T98G^Empty^ and T98G^cGAS^ cells were seeded in 150 mm dishes 18 h prior dsDNA transfection. 6 h post transfection Opti-MEM was replaced with 13.5 mL of DMEM media. Conditioned media was collected 24 h post transfection, centrifugated and filtered using 2 µm filters and frozen at -80°C.

### THP-1 polarization assay

THP-1 cell lines were differentiated in macrophages (M0) using PMA (Phorbol 12-myristate 13-acetate) at 150 nM during 24 h. Then, the media was aspirated and replaced by RPMI. Forty-eight hours later PMA-treated THP-1 were incubated with 2/3 of conditioned media complemented with 1/3 of fresh DMEM for 24 h. Cells were then harvested and samples analysed by RT-qPCR.

### Human Blood-derived cells

Buffy coats from healthy donors were obtained from the Etablissement Français du Sang (EFS, Montpellier, France). Isolation and differentiation of human CD14^+^ monocyte were performed according to previously reported protocols ^51, 52^. Briefly, freshly isolated CD14^+^ monocytes were incubated with indicated conditioned media for 72 h. Cells were then harvested and samples processed for flow cytometry. As control, CD14^+^ monocytes were also incubated for 3 days in complete media prior to analysis of polarization status by flow cytometry analysis.

### Flow cytometry

Cells (10^5^ cells/staining) were harvested and fixed in 1% paraformaldehyde for 15 minutes. Cells were then washed in PBS/1% BSA buffer and incubated for 30 minutes on ice with the following antibodies from Biolegend (San Diego, USA): FITC-anti-human CD38 (#303503), PE-anti-human CD206 (#321105), PE-anti-human CD209 (#330106), APC-anti-human CD86 (#305411). After washes in PBS/1% BSA buffer, cells were acquired on a Novocyte flow cytometer (Agilent Technologies). For blood-derived monocytes phenotyping, FITC-anti-human CD14 (Miltenyi, #130-110-576) antibody from Biolegend was also used.

### Mouse xenograft tumor model

Athymic Nude Foxn1-nu males (Envigo) of 7 weeks were sub-cutaneously injected with 5 × 10^6^ T98G^Empty^ or T98G^cGAS^ in the right or left flank, respectively. Cells were injected in a total volume of 100 µl in a mix of DMEM/Matrigel (v/v). Subcutaneous tumor growth was monitored over time and tumor size measured with a caliper. Tumor volume was estimated using the formula: volume = length × width^2^ × 0.526.

### THP-1 migration assay

24-well cell culture inserts 3.0 µm PET clear (Cell QART) were placed into the wells of a 24-well plate, containing 500 µL of conditioned media. DMEM containing FBS was used as a negative control, while DMEM with 30 ng/mL of CCL2 as a positive control. THP-1 cells were counted, resuspended at a concentration of 3 × 10^5^ cells/mL in RPMI without FBS and 100 µL of cell suspension added to the insert. Six hours later, the upper surface of the transwell membrane was washed twice with cold PBS and non-migrated cells were gently scrapped with a cotton swab. Remainder cells were fixed in 100% methanol for 10 min and nuclei stained with 4′,6-diamidino-2-phenylindole (DAPI). Transwell inserts were mounted on coverslips. Apotome Z3 microscope (Zeiss) was used to visualize and count the cells that migrate through the insert.

### Proteome Profiler

To assess differential chemokine expression, Proteome Profiler Human Chemokine Array Kit (ARY017) was used following manufacturer’s instructions. Conditioned media from T98G^Empty^ and T98G^cGAS^ cells were used.

### Immunofluorescence and microscopy analysis

Cells were seeded on glass coverslips 18 h prior to dsDNA or CPT treatment and fixed either in methanol or with 4% PFA in 2% sucrose PBS. PFA fixation was followed by permeabilization in 0.1% Triton X-100 in PBS for 5 min at room temperature (RT). After blocking in PBS containing 0.1% Tween (PBS-T) and 5% BSA for 30 min at RT, cells were incubated overnight at 4°C with dilution in PBS-T, 5% BSA. Primary antibodies used in immunofluorescence include: anti-dsDNA (ab27156, Abcam, 1:100), anti-53BP1 (MAB 3802, Sigma-Aldrich, 1:300), anti-pDNA-PKcs Ser2056 (ab18192, Abcam,1:200), anti-DNA-PKcs (ab32566, Abcam, 1:100). Secondary antibody incubation was performed for 1 hour at RT. Secondary antibodies used are: Alexa Fluor 488 goat anti-Mouse IgG, (#A11001, Thermofischer) and Alexa Fluor 594-coupled goat anti-Rabbit IgG (#R37117, Thermofischer). Nuclei were stained with DAPI and coverslips mounted in Vectashield mounting media. Images were acquired by Apotome Z3 microscope by Zeiss with ZEN (blue edition) software with a 63X oil objective, and images were processed with Omero or Fiji.

### Spheroids

T98G cells lines were prepared in DMEM media and 600 cells were distributed per well of 96 well plates (Corning ultralow attachment) prior to centrifugation for 2 min at 200g. Cells were kept in the incubator during 72 h to allow spheroid formation prior to daily scanning using a Celigo cell imaging cytometer (Nexcelom) apparatus until day 8. Volume was estimated using Celigo software.

### Zebrafish experiments

The homozygous transgenic zebrafish (*Danio rerio*) line *tg(mfap4:RFP)* was raised and maintained at Azelead’s animal facility under standard conditions. Incross of this transgenic line was performed to obtain the embryos used in the experiments. Larvae were maintained at 28 °C at a maximum density of 50 larvae per Petri dish in fish water, containing methylene blue. After 24 h, larvae were placed in fish water, containing 200 μM 1-phenyl 2-thiourea (PTU), to prevent melanin synthesis.

Transplantation needles were prepared from borosilicate glass capillaries (Outer Diameter: 1 mm, Inner Diameter: 0.75 mm, 10 cm length, Sutter Instrument), lacking the internal filament. They were pulled on a glass micropipette puller (model P-30, Sutter Instrument). The tip of the needle was broken off under a stereomicroscope (Leica M80 Stereo zoom microscope) using smooth forceps. Using microloader tips, the transplantation needle was filled with 10 µL of cell suspension. The needle was next inserted in a needle holder, that is mounted on a manipulator (Narishige) and connected to the oil manual microinjector (Eppendorf CellTram vario). After anesthesia, larvae were placed in PBS containing Tricaïne, aligned and positioned dorsally between smooth forceps and 50-100 cells were transplanted in the brain region of each embryo. Transplanted larvae were next maintained at 33°C in fish water, containing PTU. Xenotransplantation was conducted in 3 dpf embryos using T98G-GFP^Empty^ and T98G-GFP^cGAS^. Control embryos were injected with PBS. Xenotransplanted fish were used for either gene expression analysis or imaging. Briefly, for gene expression analysis, 24 h post transplantation, anesthetized embryos were manually transected with a sterile blade under a stereomicroscope to isolate the head region. 25 embryos heads were then pooled in a tube (#72.693.465, Sarstedt) containing trizol and beads (2.4 mm beads, VWR) and frozen at -80°C. Prior RNA extraction, heads were dissociated in trizol using Fastprep 24^TM^ (MP) machine and RNA extracted. The relative expression of M1 and M2 zebrafish macrophage markers were normalized to *ef1a*. For image acquisition embryos were placed individually in wells of a 96-well plate-adapted molds (Azelead), anesthetized using 0.16 mg/ml PBS/Tricaine (MS-222) and imaged from the day of transplantation (D0) up to D2. Z-stack Images were acquired through a Cell Discoverer 7 system (Zeiss).

### Analysis of zebrafish images

To estimate tumor mass, Z-projections were performed using the Zen software (ZEISS). A threshold was then applied on the Fiji software for each embryo at D0, D1 and D2 to determine tumor area. Tumor area was next normalized to the day of transplantation (D0) to evaluate tumor growth of the injected cell lines. Macrophage recruitment around the tumor site was estimated by manual counting on Fiji, by moving the focus in all Z stacks. To characterize invasiveness, the elongated cells were counted manually on Fiji. Images were prepared for publication using the Fiji program by setting the same parameters in all compared images.

### Glioblastoma RNA seq Data Retrieval and correlation analysis

For mRNA expression analysis, RNA seq data of CGGA-GBM dataset, containing 224 glioblastoma samples was used. The dataset was accessed from the GlioVis web server (http://gliovis.bioinfo.cnio.es/) and mRNA levels retrieved using the correlation tool. After plotting the mRNA level of *PRKDC* against *MB21D1* we applied a filter cut-off to determine low and high expression allowing the definition of 4 populations: *PRKDC*^low^/*MB21D1*^low^; *PRKDC*^low^/*MB21D1*^high^; *PRKDC*^high^/*MB21D1*^low^; *PRKDC*^high^/*MB21D1*^high^. mRNA levels of genes of interest were plotted based on these populations. One-way Anova with Tukey’s multiple comparisons test was used to compare gene expression among populations in glioblastoma dataset.

### Statistical analysis

Statistical analysis was performed using GraphPad Prism version 7. For statistical analysis of *in vitro* experiments, unpaired Student’s t-test was performed as indicated in figure legends. One-way Anova with Tukey’s multiple comparisons test was used to analyze gene expression amongst populations in glioblastoma datasets. One-way Anova was used for mice studies. Mann-Withney test was performed to analyze tumor growth and macrophage recruitment in zebrafish. The number of replicates in each experiment (including number of mice or zebrafish) are indicated in the figure legends. All data are expressed as mean ± SEM. The statistical parameters can be found in the figures and the figure legends. Ns: non-significant. * p < 0.05, ** p < 0.01, *** p < 0.001 and **** p < 0.0001.

