## Supplementary Material for "The crosstalk between DNA-PK and cGAS drives tumor immunogenicity"

Supplementary Figures 1-5

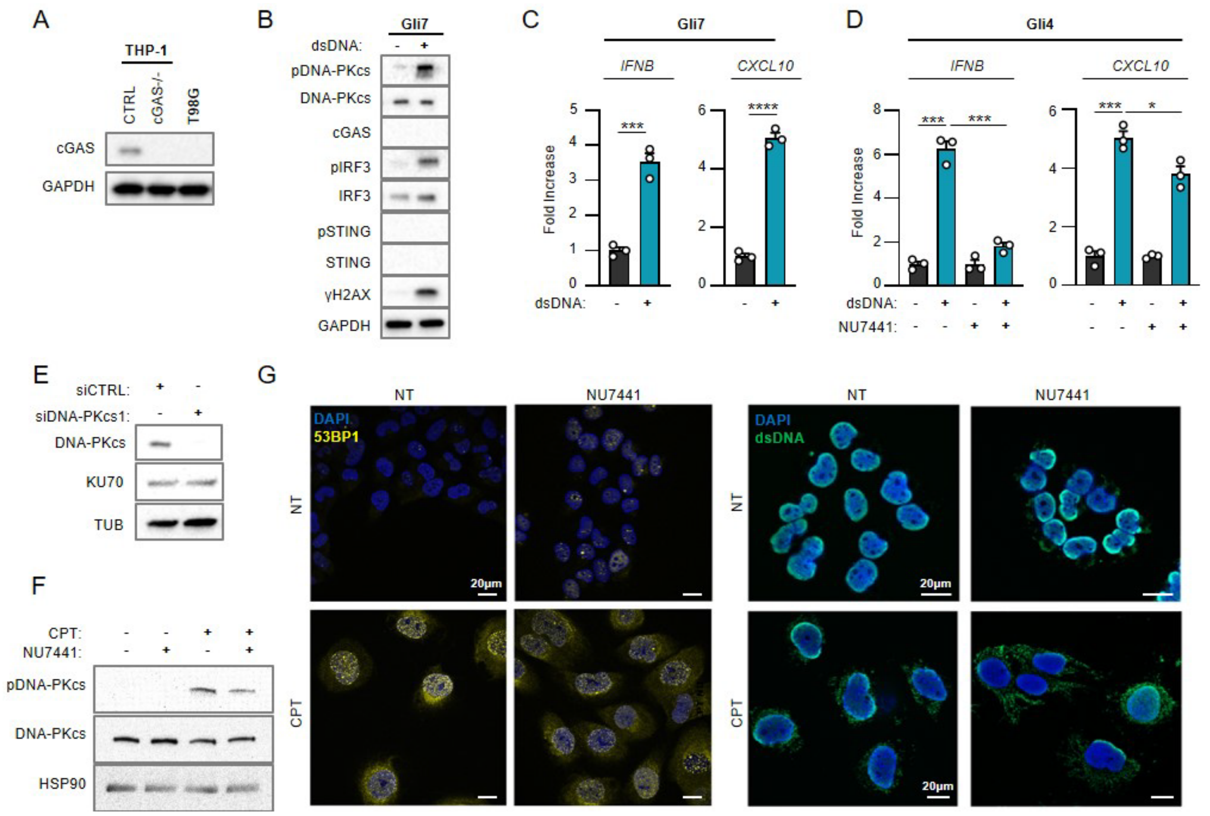


**Figure S1: DNA-PKcs promotes type I Interferon responses in cancer cells that do not express cGAS.**

(A) Whole cell extracts from THP-1$^{CTRL}$, THP-1$^{cGAS-/-}$ and T98G cells were analyzed by WB using indicated antibodies.

(B) Gli7 cells were challenged or not with dsDNA for 6 h prior to WB analysis using indicated antibodies.

(C) *IFNB* and *CXCL10* mRNA levels were analyzed by RT-qPCR in samples treated as in B (n=3).

(D) Gli4 cells were challenged or not for 6 h with dsDNA, in the presence or not of the NU7441 DNA-PKcs inhibitor, prior to analysis of *IFNB* and *CXCL10* levels by RT-qPCR. Graphs present a representative biological triplicate (n=3).

(E) T98G cells were treated with non-targeting (CTRL) or DNA-PKcs-targeting siRNAs prior to whole cell extraction and WB analysis using indicated antibodies.

(F) Whole cell extracts from T98G cells, treated or not with camptothecin (CPT) for 48 h, in presence or not of NU7441 inhibitor (24 h), were analyzed by WB using indicated antibodies.

(G) T98G cells were treated or not with CPT for 48 h, in presence or not of NU7441 inhibitor (24 h), prior to immunofluorescence analysis using dsDNA- and 53BP1-specific antibodies and DAPI nuclear staining.

All immunoblots show representative experiments. All graphs present means ± standard error from the mean (SEM). P-values were determined by Student’s t-test. * $p < 0.05$, *** $p < 0.001$, **** $p < 0.0001$.

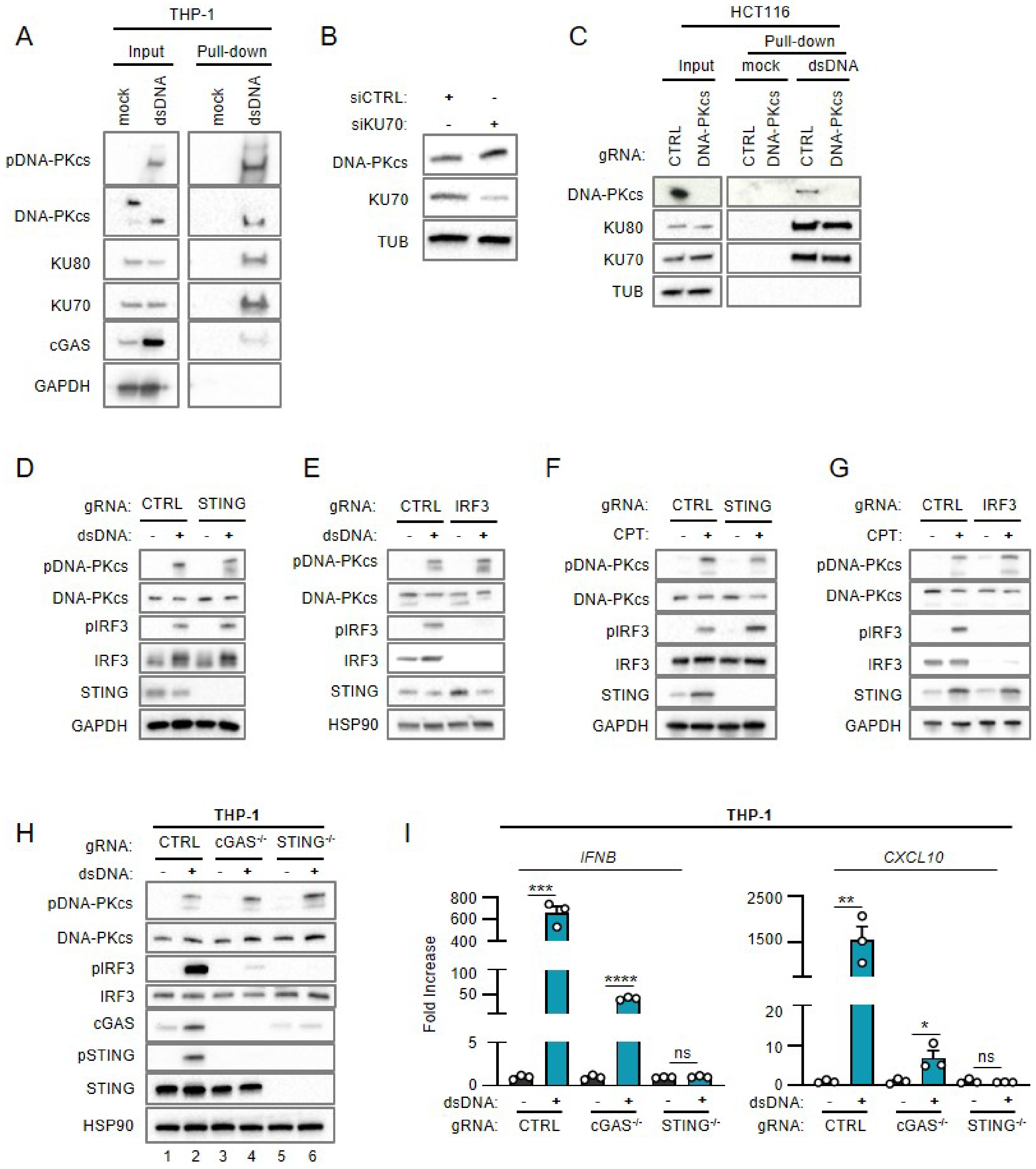


**Figure S2: DNA-PK-dependent detection of cytosolic dsDNA drives STING-independent IRF3-dependent type I IFN responses in cancer cells lacking cGAS.**
(A) THP-1 cells were transfected or not with 80nt-long biotinylated dsDNA prior to pull-down using streptavidin-affinity beads. Input and eluates were analyzed by WB using indicated antibodies.
(B) T98G cells were treated with non-targeting (CTRL) or KU70-targeting siRNAs prior to whole cell extraction and analysis by WB using indicated antibodies.
(C) Whole cell extracts from control HCT116 (CTRL) and HCT116$^{PRKDC-/-}$ were used in pull-down experiments using biotinylated dsDNA and streptavidin affinity beads. Inputs and eluates were analyzed by WB using indicated antibodies.
(D) Whole cell extracts from CTRL or *STING*-/- T98G cells transfected or not with dsDNA were analyzed by WB using indicated antibodies

(E) Whole cell extracts from CTRL or *IRF3*-/- T98G cells transfected or not with dsDNA were analyzed by WB using indicated antibodies.
(F) Whole cell extracts from CTRL or *STING*-/- T98G cells treated or not with CPT for 72 h were analyzed by WB using indicated antibodies.
(G) Whole cell extracts from CTRL or *IRF3*-/- T98G cells treated or not with CPT for 72 h were analyzed by WB using indicated antibodies.
(H) THP-1$^{CTRL}$, THP-1$^{cGAS-/-}$ and THP-1$^{STING-/-}$ were challenged or not with dsDNA for 6 h, prior to WB analysis using indicated antibodies.
(I) *IFNB* and *CXCL10* mRNA levels were analyzed by RT-qPCR in samples treated as in E. Graphs present a representative biological triplicate (n=3).
All immunoblots are representative experiments. All graphs present means ± SEM. P-values were determined by Student's t-test. ns: not significant. * $p < 0.05$, ** $p < 0.01$, *** $p < 0.001$, **** $p < 0.0001$.
Related to Figure 2.

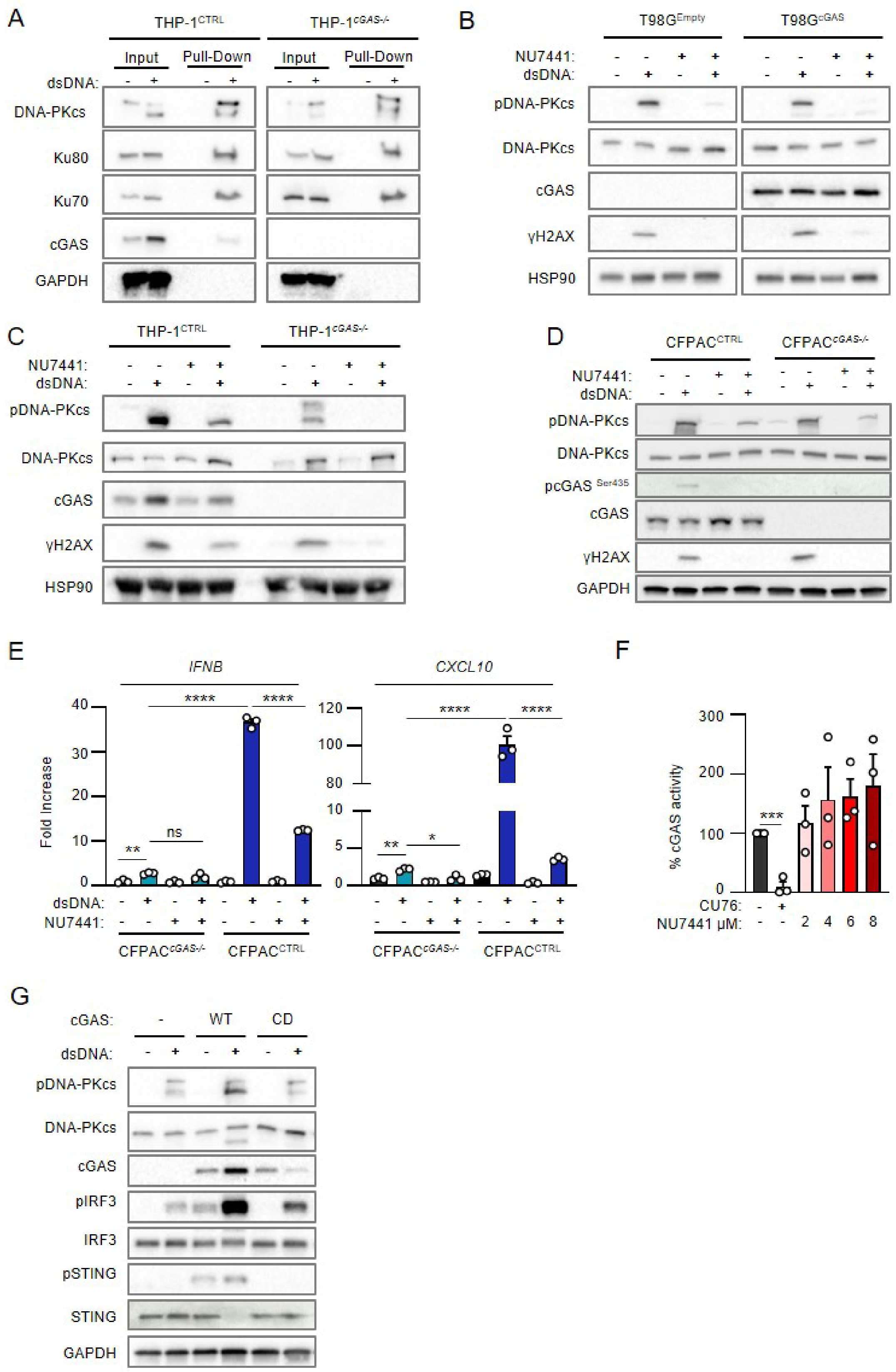


**Figure S3: DNA-PKcs and cGAS synergize for the activation of type I Interferon responses.**

(A) THP-1 and THP-1$^{cGAS-/-}$ were transfected or not with 80-nt long biotinylated dsDNA prior to whole cell extraction and pulldown using streptavidin-affinity beads. Inputs and eluates were analyzed by WB using indicated antibodies.
(B) T98G cells expressing an empty (T98G$^{\mathrm{Empty}}$) or cGAS-encoding vector (T98G$^{\mathrm{cGAS}}$) were transfected or not with dsDNA for 6 h in the presence or not of the NU7441 DNA-PKcs inhibitor. Whole cell extracts were analyzed by WB using indicated antibodies.
(C) THP-1$^{\mathrm{CTRL}}$ and THP-1$^{cGAS-/-}$ were transfected or not with dsDNA for 6 h in presence or not of the NU7441 DNA-PKcs inhibitor prior to analysis of protein expression by WB using indicated antibodies.
(D) CFPAC and CFPAC$^{cGAS-/-}$ were transfected or not with dsDNA for 6 h in presence or not of the NU7441 DNA-PKcs inhibitor prior to WB analysis using indicated antibodies.
(E) *IFNB* and *CXCL10* mRNA levels were assessed by RT-qPCR in CFPAC$^{\mathrm{CTRL}}$ and CFPAC$^{cGAS-/-}$ treated as in D. Graphs present a representative biological triplicate (n=3).
(F) cGAS activity upon treatment with 2.5 µM of the CU76 cGAS inhibitor and 2, 4, 6, or 8 µM of NU7441 was measured by ELISA (n=3).
All immunoblots show representative experiments. P-values were determined by Student's t-test. ns: not significant. * $p < 0.05$, ** $p < 0.01$, *** $p < 0.001$, **** $p < 0.0001$.

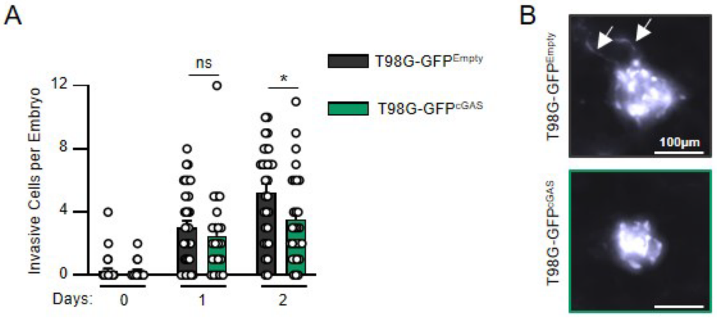


**Figure S4: Re-expression of cGAS in glioblastoma cells decrease invasiveness *in vivo*.**

(A) The number of invasive cells per embryo transplanted in Figure 4G was manually counted at D0, D1 and D2 post transplantation. Data represents mean (± SEM) of n = 21 (T98G-GFP$^{Empty}$) or 29 (T98G-GFP$^{cGAS}$) embryos. A Mann-Whitney test was performed to assess the significance * $p < 0.05$.

(B) Representative images of elongated T98G cells counted in A.

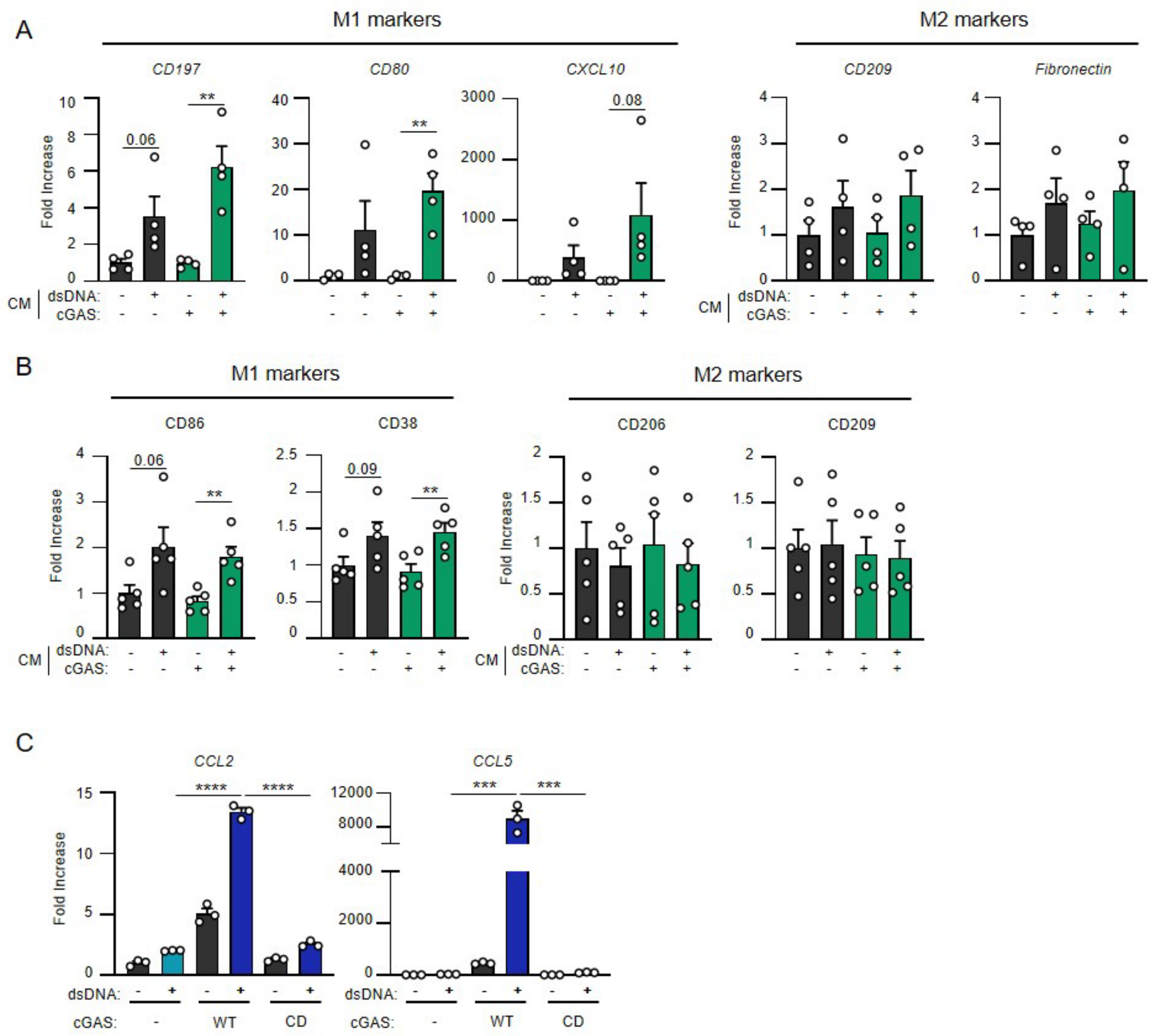


**Figure S5: Re-expression of cGAS in glioblastoma tumors promotes the secretion of chemokines that enhance macrophage recruitment.**

(A) THP-1 monocytes were incubated for 72 h with conditioned media (CM) derived from T98G$^{Empty}$ and T98G$^{cGAS}$ cells, prior to analyses of *CD197*, *CD80*, *CXCL10*, *CD209* and *Fibronectin* gene expression, by RT-qPCR (n=4).

(B) CD14+ monocytes derived from healthy donors were incubated for 72 h with conditioned media derived from T98G$^{Empty}$ and T98G$^{cGAS}$ cells, prior to flow cytometry analysis of M1 (CD86, CD38) and M2 (CD206, CD209) polarization markers (n=5).

(C) T98G$^{Empty}$, T98G$^{cGAS}$ and T98G$^{cGAS\text{-}CD}$ cells were transfected or not with dsDNA for 6 h, prior to analyses of *CCL2* and *CCL5* mRNA levels by RT-qPCR. Graphs present a representative biological triplicate (n=3).

P-values were determined by Student’s t-test. ns: not significant. ** p < 0.01, **** p < 0.0001.
